# Dense Bicoid Hubs Accentuate Binding along the Morphogen Gradient

**DOI:** 10.1101/133124

**Authors:** Mustafa Mir, Armando Reimer, Jenna E. Haines, Xiao-Yong Li, Michael Stadler, Hernan Garcia, Michael B. Eisen, Xavier Darzacq

## Abstract

Morphogen gradients direct the spatial patterning of developing embryos, however, the mechanisms by which these gradients are interpreted remain elusive. Here we perform *in vivo* single molecule imaging in early *Drosophila melanogaster* embryos of the transcription factor Bicoid that forms a gradient and initiates patterning along the anteroposterior axis. We observe that Bicoid binds to DNA with a rapid off-rate, such that its average occupancy at target loci is on-rate dependent, a property required for concentration-sensitive regulation. Surprisingly, we also observe abundant specific DNA binding in posterior nuclei, where Bicoid levels are vanishingly low. Live embryo imaging reveals spatiotemporal “hubs” of local high Bicoid concentration that are dependent on the ubiquitous maternal factor Zelda. We propose that localized modulation of transcription factor on-rates via clustering, provides a general mechanism to facilitate binding to low-affinity targets, and that this may be a prevalent feature directing other developmental transcription networks.

## INTRODUCTION

Spatial patterning during embryonic development is orchestrated through concentration gradients of regulatory molecules known as morphogens (Turing, 1952; Wolpert, 1969). The maternally deposited transcription factor (TF) Bicoid (BCD) in *Drosophila melanogaster* was the first identified morphogen (Driever & Nusslein-Volhard, 1988b), and remains an iconic and widely studied developmental regulator. Bicoid is distributed in an exponentially decaying concentration gradient along the anteroposterior (A-P) axis of embryos and predominantly regulates the activity of ~100 genes in distinct spatial expression domains ranging from the anterior tip to the middle of the embryo (Driever & Nusslein-Volhard, 1988a; Driever & Nussleinvolhard, 1989; Driever, Siegel, & Nusslein-Volhard, 1990; Struhl, Struhl, & Macdonald, 1989). The ability of BCD and other morphogens to activate target genes in different locations across gradients is classically thought to arise from the modulation of the number and strength of cognate DNA binding sites within target enhancers (Burz, Rivera-Pomar, Jackle, & Hanes, 1998; Lebrecht et al., 2005; H. Xu, Sepulveda, Figard, Sokac, & Golding, 2015), with sharp expression domain boundaries set through cooperative binding (Ephrussi & St Johnston, 2004; Lebrecht et al., 2005).

In recent years this classical model of concentration dependent activation has been challenged through experiments on mutant embryos with flattened BCD distributions which reveal that segment order and polarity can be maintained even without a concentration gradient (Ochoa-Espinosa, Yu, Tsirigos, Struffi, & Small, 2009). It has been suggested that instead of a pure concentration dependence, the activation of BCD target genes and the resulting sharp expression domain boundaries is tightly regulated by spatially opposing gradients of repressors (Chen, Xu, Mei, Yu, & Small, 2012) and the combinatorial actions of other transcription factors (P. Combs & Eisen, 2017). The recent discovery of the ubiquitous factor Zelda and its role in the regulation of chromatin accessibility (Foo et al., 2014; Harrison, Li, Kaplan, Botchan, & Eisen, 2011; Li, Harrison, Villata, Kaplan, & Eisen, 2014; Liang et al., 2008; Schulz et al., 2015; Sun et al., 2015; Z. Xu et al., 2014) and in modulating the timing and strength of Bicoid (Hannon, Blythe, & Wieschaus, 2017; Z. Xu et al., 2014) and Dorsal (Foo et al., 2014) controlled enhancer activation in a concentration dependent fashion has further strengthened the hypothesis (Lucchetta, Lee, Fu, Patel, & Ismagilov, 2005) that the interpretation of the BCD and other morphogen gradients is more complex than previously thought.

The validity of these models and essential mechanistic questions about how BCD, and morphogens in general, differentially activate genes along a concentration gradient have been challenging to resolve with current approaches. For example, extant models cannot address how BCD is sufficient for activating its targets such as Knirps (Rivera-Pomar, Lu, Perrimon, Taubert, & Jackle, 1995) and Hairy (La Rosee, Hader, Taubert, Rivera-Pomar, & Jackle, 1997), in the posterior of the embryo, where BCD nuclear concentrations are <2nM (Morrison, Scheeler, Dubuis, & Gregor, 2012) in the short interphase times (5-10 minutes) of the early nuclear cycles. Since the question of how BCD molecules can find their targets in these short times requires dynamic measurements, genomic assays and biochemical approaches which provide static snapshots are inadequate. In this study, we address this gap in our understanding by performing direct measurements of BCD-DNA interactions *in vivo* by single molecule imaging.

Single molecule imaging in living cells has been increasingly used in recent years to measure the dynamics of TF-DNA interactions (Liu, Lavis, & Betzig, 2015). However, the techniques commonly used are not suitable for whole embryos and thick tissues. Total Internal Reflection (TIRF) and Highly-Inclined illumination (Hi-Lo) (Tokunaga, Imamoto, & Sakata-Sogawa, 2008) which have enabled single molecule imaging in monolayer cell cultures use wide-field excitation geometries and restrict the illumination volume to a small distance above the microscope coverslip in order to limit the excitation of out-of-focus fluorophores. This confinement of the illumination volume is necessary to achieve the signal-to-background ratios (SBRs) required for single molecule detection. Consequentially, if the thickness of the illumination volume is extended to image further away from the coverslip, the SBR degrades as more and more out-of-focus fluorophore emission raises the background level and reduces contrast. This degradation is further exacerbated when imaging highly auto- fluorescent samples such as embryos or thick tissues.

Lattice Light-Sheet Microscopy (LLSM) was recently developed to overcome these technical barriers (B. C. Chen et al., 2014). Here we apply LLSM to developing *Drosophila melanogaster* embryos in order to characterize the single molecule DNA-binding kinetics of BCD. We find that BCD binds to chromatin in a highly transient manner across the concentration gradient, and were surprised to observe a significant number of binding events even at low concentrations in the posterior-most regions of the embryo. Examination of the spatial distribution of BCD binding events reveals spatiotemporal hubs of high local BCD concentration. Through genome wide analysis of BCD-DNA binding on dissected posterior segments of embryos, we show that the binding we observe via single molecule imaging in posterior nuclei occurs at specific regulatory regions. We find that the regions which are enriched for BCD in the posterior segments are highly correlated with Zelda (ZLD) binding, signifying its role in mediating the formation of BCD hubs. Through single molecule imaging of BCD in ZLD null embryos we show that ZLD is required for the formation of BCD hubs in the posterior embryo. Together, these data advocate for a model in which ZLD mediates the formation of hubs of high local BCD concentration and enables the activation of BCD dependent targets at all position across the anteroposterior axis of the embryo.

## RESULTS

### Single molecule imaging in living Drosophila embryos using Lattice Light-Sheet Microscopy

The principle of LLSM (Figure 1-figure supplement 1) is to create an excitation light-sheet that matches the depth-of-field of the detection objective such that only fluorophores that are in focus are excited (B. C. Chen et al., 2014). As in all light-sheet microscopes, in LLSM the excitation and detection objectives are independent and oriented orthogonally to each other. However, unlike conventional light-sheet modalities that use Gaussian beam illumination, in LLSM an array of Bessel beams is used. The spacing and phase of the Bessel beams in the array is controlled such that their side-lobes destructively interfere in order to achieve maximal axial confinement of the light-sheet. Thus, unlike in wide-field excitation geometries the thickness of the excitation volume in LLSM is independent of the distance from the coverslip which is being probed. This approach enables single molecule imaging of BCD-eGFP in living *Drosophila* embryos over a large field of view with high temporal resolution (Figure 1 A-B, Figure 1-figure supplement 2 and Videos 1-3). We utilize a *yw*; *his2av-mrfp1; bcdE1, egfp-bcd* fly line in which only the labelled BCD is expressed indicating proper functionality and expression levels (Gregor, Wieschaus, McGregor, Bialek, & Tank, 2007) to ensure that all molecules we observe are functionally relevant. The single molecule nature of the data is reflected in the distribution of intensities (Video 2 and Figure 1-figure supplement 2A-B) and discrete characteristics (Video 2 and Figure 1-figure supplement 2C-D) of the observed binding events.

**Figure. 1.**
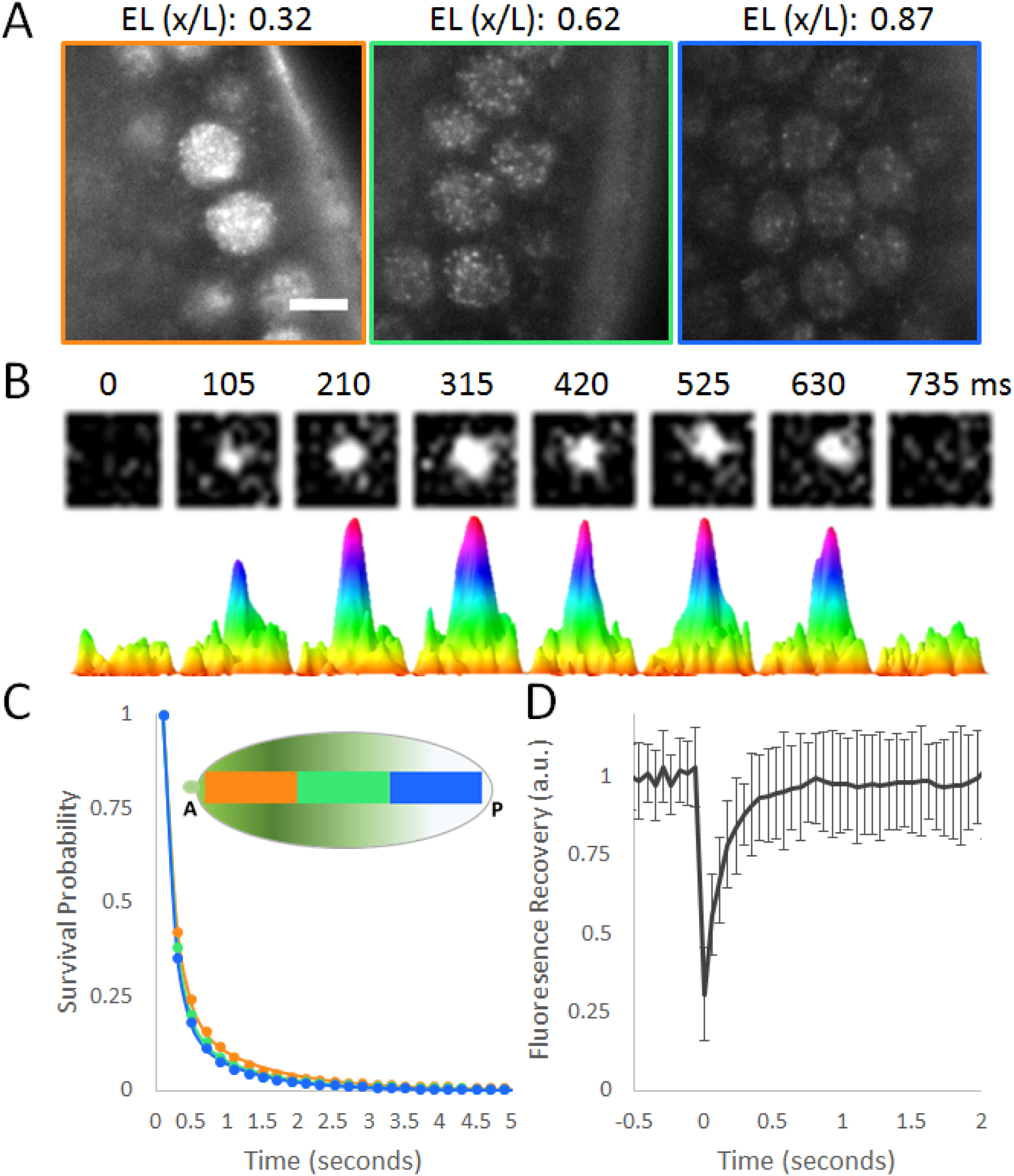
Single molecule kinetics of Bicoid in living *Drosophila* Embryos. **(A)** Raw images of BCD-eGFP molecules in a living *Drosophila* embryo acquired with a 100 millisecond exposure time. Scale bar is 5 um. Positions along the A-P axis are shown as a fraction of the Embryonic Length (EL, x/L). **(B)** Example of a single molecule binding event. Top row shows raw images from a 1.2 x 1.2 μm area, bottom row shows corresponding surface plot representations to illustrate the signal-to-noise. **(C)** Uncorrected survival probability curves for Bicoid binding (markers) in the Anterior (34 nuclei), Middle (70 nuclei) and Posterior (83 nuclei) segments of the embryo and corresponding fits to a two-exponent model (solid lines) show no significant differences. **(D)** FRAP curve for Bicoid shows a recovery time on the order of hundreds of milliseconds, error bars show standard deviation over 21 nuclei.

Bicoid nuclear concentrations range from ~50 nM at the anterior most positions down to <2 nM (Morrison et al., 2012) in the posterior. This translates to a range on the order of 10^4^-10^2^ BCD molecules/nucleus. Assuming an isotropic distribution of molecules and a 400 nm thick imaging volume, we can estimate a range on the order of 10^3^-10^1^ BCD molecules/ imaging plane. This range of concentrations is reflected in the data shown in Video 1, where unambiguous single molecule detections can be seen in the middle and posterior positions from the start, whereas in the anterior positions they can only be detected when a sufficient amount of bleaching has occurred. This natural concentration range allows us to perform single molecule tracking at all positions in the embryo without utilizing sparse labelling strategies or photo-switchable fluorophores.

### Bicoid binds chromatin in a highly transient manner across the concentration gradient

The classical model for morphogens predicts a difference in the average dissociation rates of Bicoid along the A-P axis. For example, genes that are activated at lower concentrations should have higher affinity sites resulting in lower off-rates to enable a higher time-average occupancy (i.e. residence time, RT). To test this model at the single molecule level we therefore first performed single molecule imaging and tracking at long (100 millisecond) exposure times, effectively blurring out the fast moving (unbound) population (Videos 1-2 and Figure 1-figure supplement 2) (J. J. Chen et al., 2014) to estimate the residence times of BCD binding in nuclei at all positions along the A-P axis.

Previous single molecule studies of transcription factors have consistently found two populations in the survival probability distributions, a short-lived population with RTs on the order of hundreds of milliseconds, and a longer-lived population with RTs on the order of 10’s of seconds to minutes (J. J. Chen et al., 2014; Hansen, Pustova, Cattoglio, Tjian, & Darzacq, 2017; Normanno et al., 2015). These two populations have often been shown to be the non-specific and specific binding populations, respectively. The survival probability distributions of BCD similarly are fit better with a two-exponent model than with a single-exponent model indicating the presence of two sub-populations (Figure 1-figure supplement 3 and Figure 1-figure supplement 4). Fits to the survival probability distributions of BCD binding events (Figure 1C and Table 1) in the anterior, middle and posterior thirds of the embryo identified short-lived populations with average RTs (after photo-bleaching correction) on the order of 100s of milliseconds and longer lived populations with average RTs on the order of seconds, and with no significant dependence on position along the A-P axis for either population (Table 1). This finding is contrary to the classical model which predicts a modulation of binding affinity as the mechanism for concentration readout. The validity of our RT estimations are supported by additional measurements at 500 millisecond exposure times (Figure 1-figure supplement 4).

**Table 1.** Results from 2-exponent model fits to survival probability distributions. k_ns_ and k_s_ are the un-corrected off-rates for the short-lived (non-specific) and longer-lived (specific) populations respectively determined from a two-exponent fits to the survival probability distributions. The binding time (one-over the off rates) are shown with 95% confidence intervals in square brackets. The photo-bleaching corrected binding times are calculated as 1/(k_s_-k_bleach_) where k_bleach_ is 0.0043 s^−1^).

| Data Set | A-P Poistion | Non-specific binding time ($1/k_{ns}$ (sec)) | Specific Binding time ($1/k_s$ (sec)) | Corrected Specific Binding time $1/(k_s - k_{bleach})$ (sec) | Number of trajectories |
| --- | --- | --- | --- | --- | --- |
| 100 ms | Ant | 0.175<br>[0.166, 0.184] | 1.100<br>[1.013, 1.203] | 1.105 | 17735 |
|  | Mid | 0.160<br>[0.152, 0.168] | 0.940<br>[0.859, 1.038] | 0.944 | 40092 |
|  | Post | 0.146<br>[0.141, 0.151] | 0.865<br>[0.806, 0.932] | 0.868 | 20823 |
|  | All Together | 0.159<br>[0.166, 0.152] | 0.959<br>[0.882, 1.050] | 0.963 | 78650 |
| 100 ms (Zelda-) | Ant | 0.152<br>[0.146, 0.160] | 0.777<br>[0.726, 0.836] | 0.780 | 11415 |
|  | Mid | 0.131<br>[0.126, 0.137] | 0.629<br>[0.587, 0.679] | 0.631 | 7572 |
|  | Post | 0.112<br>[0.108, 0.117] | 0.463<br>[0.435, 0.495] | 0.464 | 3606 |
|  | All Together | 0.139<br>[0.133, 0.145] | 0.694<br>[0.647, 0.748] | 0.696 | 22593 |
| 500 ms | All Together | 0.265<br>[0.255, 0.277] | 1.876<br>[1.69, 2.107] | - | 1211 |

Remarkably, the RT of the long-lived population is much shorter than lifetimes (10-60 sec) typically observed for other sequence specific DNA binding TFs using single molecule tracking (J. J. Chen et al., 2014; Hansen et al., 2017). The transient nature of BCD binding is further supported by Fluorescence Recovery After Photobleaching (FRAP) experiments (Figure. 1D) which reveal fast halftimes of recovery on the order of hundreds of milliseconds (Figure 1- figure supplement 5) and previous measurements by others using Fluorescence Correlation Spectroscopy (Porcher et al., 2010). The dominance of the short-lived interactions (Figure 1C) highlights the preponderance of low-affinity BCD binding sites in the *Drosophila melanogaster* genome and the resulting large number of non-specific interactions as previously suggested by genomic studies that captured its promiscuous binding behavior (Li et al., 2008; Ochoa-Espinosa et al., 2005). In addition to the expected large number of binding events in the anterior and middle segments of the embryo, we also observed a surprisingly large number of potentially specific binding events in the posterior-most nuclei where BCD has been reported to be at vanishingly low (<2nM posterior vs ~50 nM anterior) concentrations (Morrison et al., 2012).

### Spatiotemporal hubs of Bicoid binding enrich local concentrations in the posterior embryo

The lack of a detectable difference in the survival probability distributions along the A-P axis and the observation of significant binding in posterior nuclei prompted us to examine how much of the small BCD population remaining in the posterior embryo is actually bound vs. mobile. Since longer exposure times only allow detection of molecules bound for at least the span of the exposure and don’t provide any data on the mobile population, we performed single molecule tracking measurements at a decreased exposure time of 10 milliseconds. The signal-to-noise ratios at these lower exposure times are adversely affected as expected, limiting the type of analysis that can performed on this data (Video 3). However, despite this reduced contrast we were able to perform single particle tracking, and through analyses of displacement distributions (Figure 2 –figure supplement 1), we estimated the fraction of BCD that is bound along the A-P axis (Figure. 2A). This analysis suggests that a greater fraction of the BCD population is bound in more posterior positions of the embryo where BCD is present at the lowest concentrations. This counter-intuitive result motivated us to further examine the 100 millisecond exposure time dataset beyond the analysis of residence times described above.

**Figure. 2.**
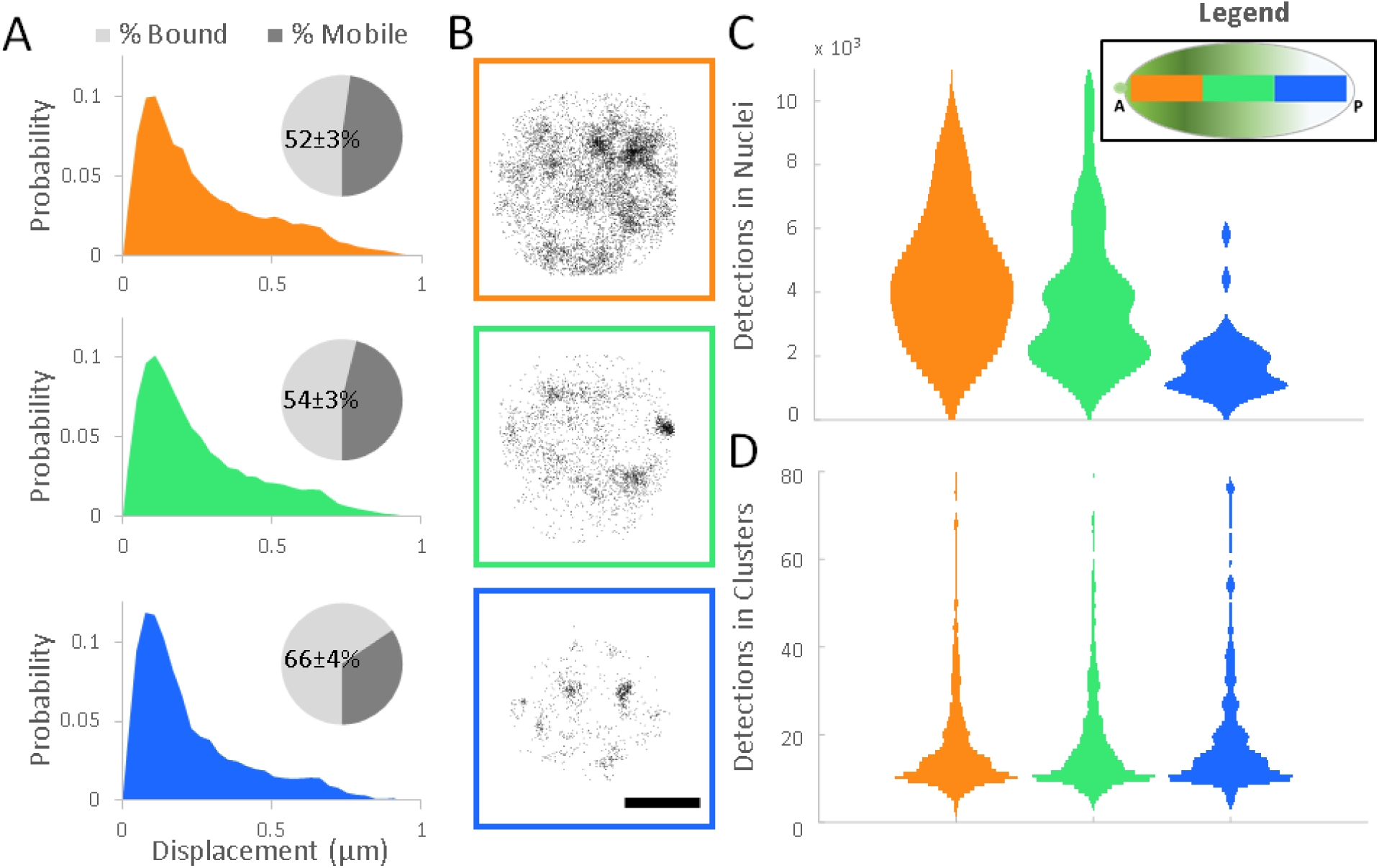
Local modulation of Bicoid Concentration. **(A)** Normalized probability distributions of measured displacements in the Anterior (30 nuclei), Middle (67 nuclei), and Posterior (66 nuclei) positions of the embryos, pie charts show the estimated mobile and bound fractions from fits to a two population distribution with the bound population percent labelled with the standard error of the fit parameter. **(B)** Examples of the spatial distribution of all detections in nuclei along the A-P axis, scale bar is 2.5 μm. **(C)** Distribution of the number of detections in all nuclei. **(D)** Distributions of the number of detections within all clusters.

Analyses of the spatial distribution of binding events in the 100 millisecond dataset revealed a distinct clustering of binding events which becomes more pronounced toward posterior positions (Figure 2B, Figure 2- figure supplement 2). Remarkably, although the number of binding events per nucleus follows the trend dictated by the global concentration gradient across the embryo (Figure 2C), the distribution of BCD molecules detected per cluster or “hub” is maintained even in the posterior (Figure 2D). These data suggest that the formation of BCD hubs with high time-averaged occupancy at specific sites in nuclei across the A-P axis is conserved independent of the global concentration gradient. This surprising finding raises the questions of whether the binding in the hubs observed in posterior nuclei is to specific targets and whether a mechanism then exists to selectively enrich local time-averaged concentrations in order to promote these interactions.

### Bicoid binds specific regulatory regions in the posterior embryo in a Zelda dependent manne

To test whether BCD is binding with specificity in the posterior embryo we analyzed its binding profiles in a spatially segregated manner (P. A. Combs & Eisen, 2013) by comparing ChIP-seq profiles derived from individually dissected posterior thirds of embryos to previously published data from whole embryos (Bradley et al., 2010). Our analysis reveals that BCD indeed binds to known targets in the posterior but with increased relative enrichment at specific enhancer elements (Figure 3A). For example, in the *hunchback* locus, binding at the posterior stripe enhancer (Perry, Bothma, Luu, & Levine, 2012) is highly enriched over the background in nuclei from the dissected posterior third relative to the whole embryo. Intriguingly, genomic regions that exhibit a relative increase in BCD occupancy in the posterior are correlated with an enrichment of Zelda (ZLD) binding (Figure 3A and Figure 3-figure supplement 1), a ubiquitous activator often described as a pioneer factor active during early embryonic development (Foo et al., 2014; Harrison et al., 2011; Liang et al., 2008; Z. Xu et al., 2014). Remarkably, enrichment of ZLD is more predictive of BCD binding in the posterior than previously determined position of enhancer activity for the loci shown in Figure 3A. Analysis of the correlation between BCD and ZLD enrichment at the cis-regulatory modules of 12 gene loci, and at ZLD and BCD peaks genome-wide reveal that binding of BCD in posterior nuclei is highly correlated with ZLD co-binding (Figure 3A, Figure 3-figure supplement 1, and Figure 3-figure supplement 2).

**Figure. 3.**
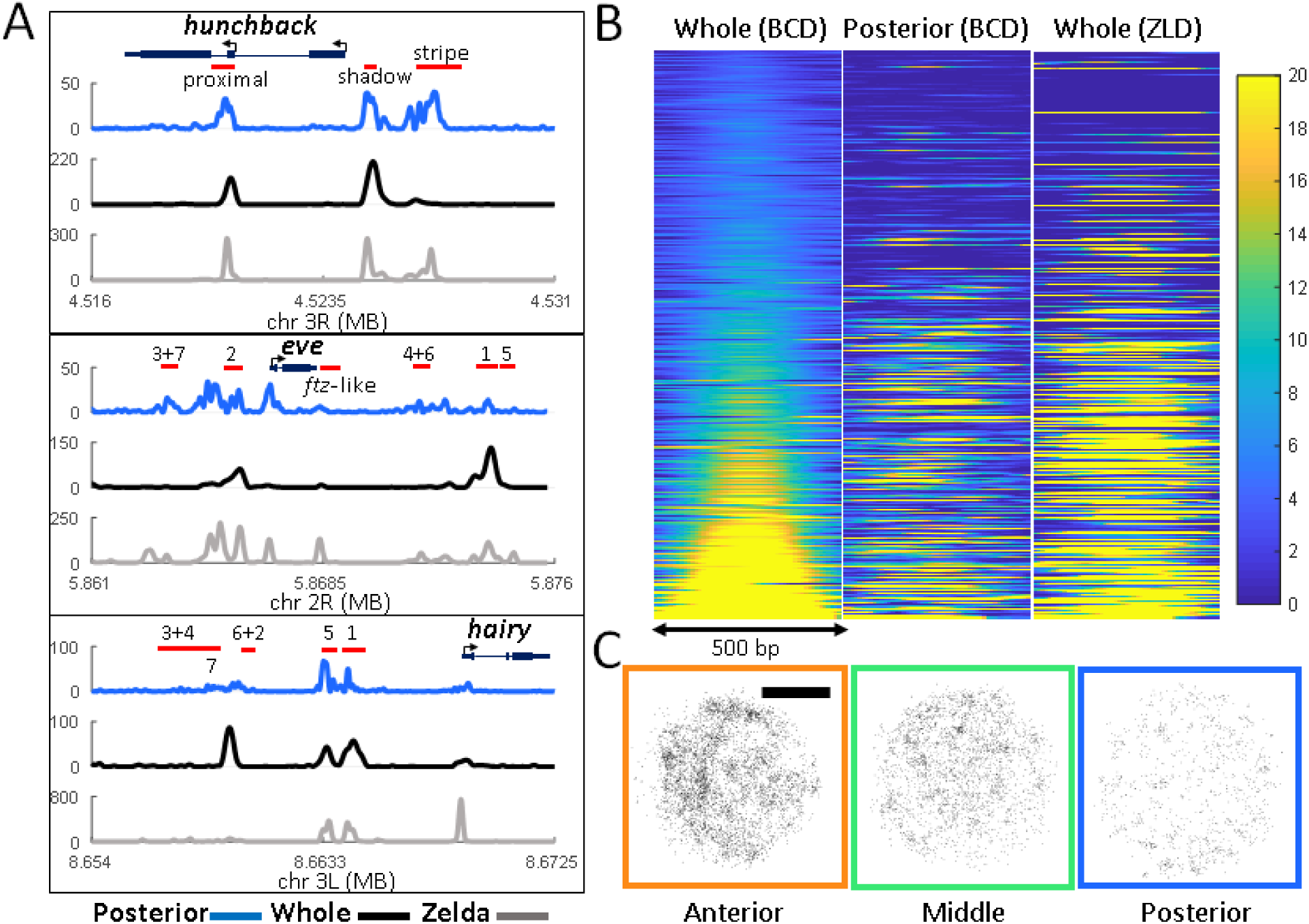
Zelda mediated Bicoid binding in the posterior embryo (A) Posterior third (blue) and whole embryo (black) BCD, and whole embryo Zelda (grey) ChIP-seq signal normalized reads at the *hunchback*, *eve*, and *hairy* gene loci. Red bars show known enhancers as annotated in the RedFly database, for eve and hairy they are numbered according to the stripes they are thought to regulate. **(B)** Heat-map representation of normalized BCD ChIP-seq reads (first two panels) and ZLD-ChIP-seq reads (third panel) in a 500 bp window centered on BCD peaks called in the whole embryo data and sorted according to increasing signal of the whole embryo data, a total of 2145 peaks are shown, colors indicate enrichment over the background (blue) with all plots displayed on the same scale. **(C)** Examples of the spatial distribution of all detected bound molecules in nuclei along the A-P axis in ZLD- embryos, scale bar is 2.5 μm. A loss of clustering is apparent compared to the distributions shown in Figure 2B.

### Formation of Bicoid hubs in the posterior embryo is dependent on Zelda

The posterior thirds genomic data and the published evidence for Zelda’s role in the regulation of chromatin accessibility (Foo et al., 2014; Harrison et al., 2011; Li et al., 2014; Liang et al., 2008; Schulz et al., 2015; Sun et al., 2015; Z. Xu et al., 2014) and its suggested role in the modulation of TF binding at low concentrations (Schulz et al., 2015; Z. Xu et al., 2014) naturally led us to hypothesize that the observed clustering of BCD binding events may be, in part, mediated by ZLD. We thus generated *zelda* null embryos and measured BCD binding at 100 millisecond exposure times, and found an abolishment of BCD hubs in the posterior embryo and a small but insignificant decrease in RTs (Figure 3C, Table 1). Due to this loss of clustering the same analysis that was performed in the WT case (Figure 2C) could not be done in the ZLD mutants. We thus calculated the pair-correlation function for the spatial distribution of binding events in both the WT and ZLD mutants (Cisse et al., 2013). This analysis allows us to infer whether binding events are spatially randomly distributed or clustered (Figure 3-figure supplement 3). Both the magnitude and correlation length indicate a diminishment of clustering in the posterior nuclei of the ZLD mutants. We also note that due to the lower labelling density of BCD in the *zelda* null embryos the presumably ZLD independent clustering in the anterior embryo now becomes more apparent (Figure 3-figure supplement 3). The loss of clustering in the ZLD mutants also confirms that the clustering we originally observed is not due to aggregation of eGFP, or other artifacts reasons.

Although the exact mechanism by which ZLD mediates BCD hub formation and binding remains unclear, we speculate that a combination of protein-protein interactions facilitated by intrinsically disordered low-complexity domains(Hamm, Bondra, & Harrison, 2015) of ZLD and its reported role in promoting chromatin accessibility (Foo et al., 2014; Li et al., 2014; Schulz et al., 2015; Sun et al., 2015) may contribute to BCD clustering (Figure 4).

**Figure. 4.**
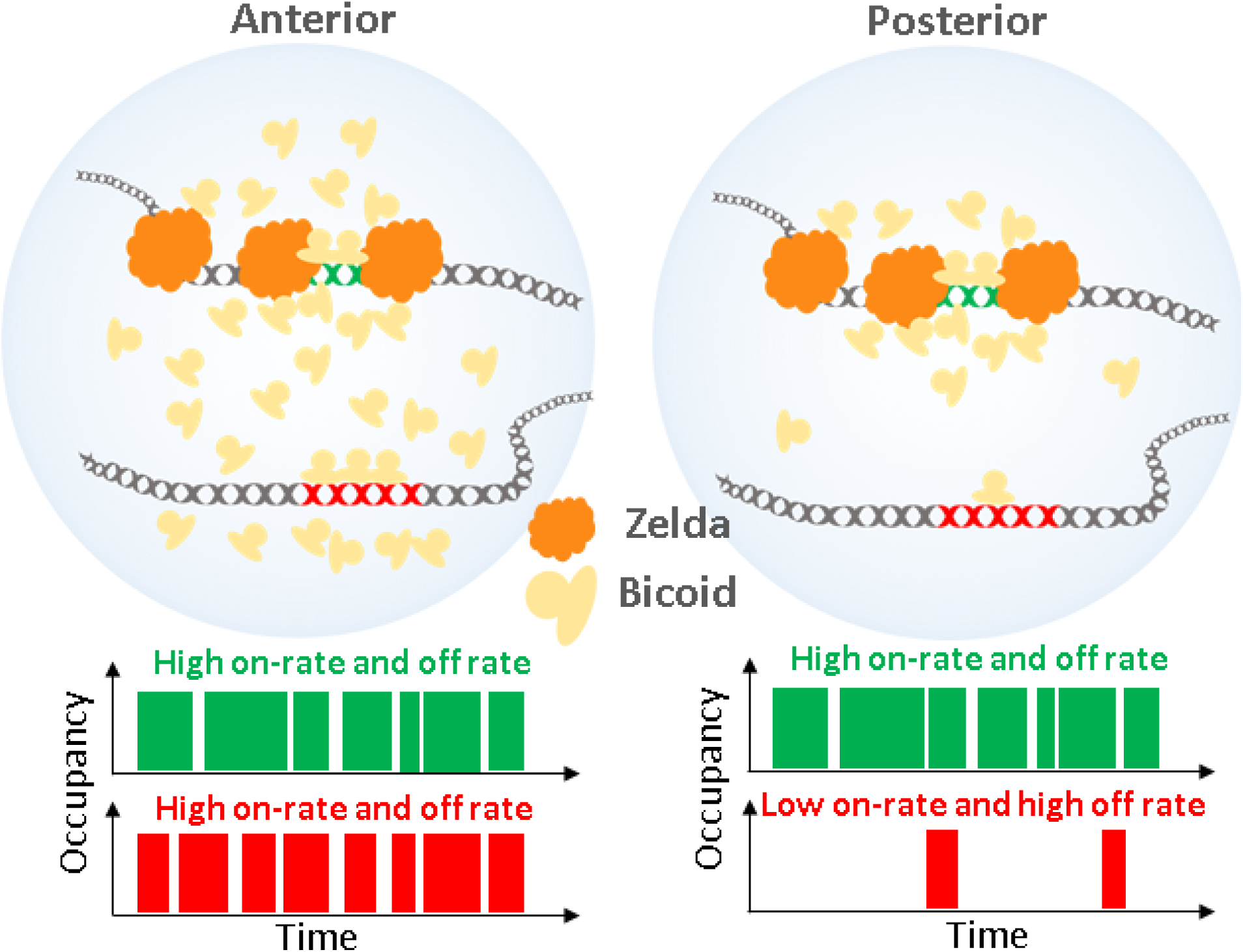
Model of Zelda dependent modulation of the Bicoid on-rate at specific loci in the posterior embryo. At high concentrations in the anterior of the embryo, all target sites are highly occupied. At low concentrations, loci with Zelda occupancy have an increased time averaged occupancy through the formation of spatiotemporal hubs that enrich local concentrations and increase the on-rate.

## DISCUSSION

Our initial observation of the low affinity nature of BCD binding to chromatin fits the classical view of BCD as a concentration-dependent morphogen. We envision that high TF concentrations (in the anterior embryo) potentiate rapid on-rates along with a high chromatin occupancy. The observed high off-rate also suggests that BCD is free to frequently sample non-specific sites (Figure 4). Thus, as BCD concentrations decrease posteriorly along the gradient, there should be progressively lower on-rates and the binding at specific sites should be correspondingly diminished, regardless of the roles of opposing repressor gradients. However, this simplistic model essentially consigns BCD to no posterior function and contradicts a wealth of evidence in the literature pointing to a specific role for BCD in the regulation of posterior gene expression (Burz et al., 1998; Chen et al., 2012; La Rosee et al., 1997; Ochoa-Espinosa et al., 2009; Rivera-Pomar et al., 1995; Small, Blair, & Levine, 1996). Although this paradox is inexplicable under the classical morphogen model and not addressed by more recent studies that cannot access dynamic information, our evidence for local high-concentration BCD “hubs” capable of modulating specific binding with help from ZLD in the posterior suggests a novel mechanism for TFs to carry out their regulatory role even when present at very low concentrations (Figure. 4). The formation of such clusters or hubs has been reported for other TFs (J. J. Chen et al., 2014; Justin Crocker et al., 2017; Liu et al., 2014) and for RNA Polymerase II (Cisse et al., 2013) and indicate that such spatial organization of the nucleus may be a general mechanism to catalyze important regulatory interactions. During embryonic development, it is likely that clustering of TFs mediated by co-factors has evolved in eukaryotes to allow exquisite spatial and segmental modulation during development through enabling interactions with low-affinity enhancers (J. Crocker, Noon, & Stern, 2016).

## MATERIALS AND METHODS

### Fly husbandry

All fly cages were prepared by combining males and females of the desired strains in a plastic cage left at room temperature in light at least 3 days prior to imaging. The lids on the cages were filled with agar dissolved in apple juice (2.4% g/w Bacto agar, 25% apple juice, 75% distilled water, and .001% of mold inhibitor from solution of 0.1g/ml (Carolina 87-6165). A paste of dry yeast was smeared on the lids to induce egg laying. Lids were exchanged once each day.

The fly strain used for all WT BCD imaging experiments was: *yw*; *his2av-mrfp1; BcdE1, egfp-bcd*. This fly line results in embryos where only the labelled BCD is expressed indicating proper functionality and expression levels (Gregor et al., 2007). For the *zld-* experiments, *bcd-egfp* heterozygous virgins with *zld-* germline cells (maternal germline clones prepared as in (Liang et al., 2008)) were crossed to *yw* males, and progeny were used for imaging 2-3 hours after laying. The heterozygosity results in only half of the BCD labelled in the *zld-* embryos. For photo-bleaching controls the line used was *yw, his2av-egfp;* +/+ (Bloomington # 24163).

### Live embryo collection for imaging

For embryo collection, lids on fly cages were exchanged one hour prior to imaging. After one hour, embryos were collected from the lids using an inoculation loop. A 5mm diameter glass cover slip (#64-0700, Warner Instruments) was prepared by immersion in a small amount of glue (prepared by dissolving adhesive from about 1/5 of a roll of double-sided Scotch tape overnight in heptane) and was left to dry for 5-10 minutes while collecting embryos. Collection was performed on a dissection scope with trans-illumination. Embryos were bathed in Halocarbon oil 27 (Sigma) for staging and then selected between developmental stages 1 and 4. Selected embryos were placed on a small square of paper towel, then de-chorionated in 100% bleach for 1 minute. Bleach was wicked off with a Kimwipe after one minute, then the square was washed with a small amount of distilled water. Excess water was wicked off the square and the square was dipped in a small water bath. Un-punctured embryos that floated to the top of the bath were selected for imaging and placed on a small paper towel square to slightly dry. To prevent excess desiccation embryos were immediately placed on the glass cover slip in rows and then immersed in a drop of phosphate-buffered saline (PBS).

### Lattice Light-Sheet Microscopy

Imaging was performed using a home built lattice light-sheet microscope (Figure 1-figure supplement 1) following the design described by Chen et al. (B. C. Chen et al., 2014) and detailed blueprints provided by the Betzig group at HHMI Janelia Research Campus. To perform the single molecule experiments we added a detection module containing two EMCCDs (Andor iXon Ultra) for dual color imaging. The EMCCDs provided a significant improvement in signal to noise over the sCMOS (Hamamatsu Orca Flash V2.0) used in the original system and made it possible to use lower excitation powers while maintaining single molecule sensitivity. In brief, the output beam from each laser is expanded and collimated independently to a size of 2.5 mm. The expanded beams for each laser are combined and input into an Acoustic Optical Tunable Filter (AOTF) to allow for rapid switching between excitation wavelengths and adjustment of power (Figure 1-figure supplement 1A). A pair of cylindrical lenses is then used to elongate and collimate the output Gaussian beam to illuminate a stripe on a spatial light modulator (SLM). The SLM is used to generate a coherent pattern of an array of 30 Bessel beams spaced such that they coherently interfere to create a 2D optical lattice pattern with a maximum numerical aperture (NA) of 0.6 and minimum NA of 0.505. A 500 mm lens is used to project the Fourier transform of the SLM plane onto an annular mask, conjugate to the back pupil plane (BPP) of the excitation objective, to spatially filter the pattern (Figure 1-figure supplement 1B). The BPP is then projected first onto a galvo scanning mirror for z-scanning and then onto a second galvo scanning mirror for x-dithering. The x-galvo scanning plane is projected onto the BPP of the excitation objective (Figure 1-figure supplement 1C). The excitation objective focuses the lattice pattern onto the sample, exciting any fluorophores within the axial range (~400nm) of the sheet. The emitted fluorescence is collected by the detection objective which is oriented orthogonally to the excitation objective and projected onto an intermediate image plane by a 500 mm tube lens (Figure 1-figure supplement 1D). An 80 mm and 200 mm lens pair is then used to de-magnify the image further to provide a 100 nm sampling per pixel on each of the EMCCD sensors. A dichroic mirror (Semrock FF560-FDi01) is placed between the last lens pair to allow for dual color imaging in red and green with maximal spectral separation. An emission filter is placed in the path of each camera to both reject the excitation wavelengths and also select the wavelength range of interest (Semrock FF03-525/50 for eGFP and FF01-593/46 for RFP) (Figure 1-figure supplement 1E). During each camera exposure the x-galvo mirror is dithered twice over a 5.1 μm range in 100 nm steps to provide uniform illumination.

The prepared coverslip, with embryos arranged in rows as described above, was then loaded into the sample holder and secured on to the positioning stages of the lattice light-sheet microscope. The sample chamber was filled with PBS for imaging and kept at room temperature. The slide was then scanned to find an embryo of suitable age (between nuclear cycles 10 and11) and the positions of the anterior and posterior extremes of the embryo were then marked. For each dataset acquired, the stage position was recorded to determine the position as a fraction of the embryonic length (EL) as the distance of the position from the anterior pole divided by the total length of the embryo.

For the residence time measurements on BCD-eGFP, a 488 nm excitation laser was used with a power of 2.9 mW measured at the back pupil plane of the excitation objective. Images were acquired with 100 millisecond exposure times and an EM Gain setting of 300. At each location at least 1000 frames were acquired resulting in a total time of 105 seconds, with a frame rate of 105 ms. Prior to and after acquiring the BCD-eGFP data an image was taken in His2-AVmRFP using a 561 nm excitation laser at an excitation power of ~0.17 mW at the back pupil plan to determine the nuclear cycle phase, data not acquired during interphase was discarded upon examination of these images. For residence time measurements at 500 ms exposure times, the 488 nm excitation laser power at the back pupil plane was reduced to 0.5 mW all other settings were the same as above. For the displacement distribution measurements the exposure time was set to 10 ms, resulting in a frame rate of 15 ms and the excitation power was increased to 8.28 mW for the 488 nm laser line. All other settings were the same as described above. Viability of the embryos was determined by allowing them to develop till gastrulation after imaging. For the *zelda*- embryos lethality was confirmed after imaging.

### Curation of data for analysis

For all datasets the following procedure was followed: First, for each movie the corresponding before-and-after histone images were checked for any evidence of chromatin condensation to ensure that analysis was only performed in interphase nuclei. Data from mid-to-late nuclear cycle 14, where the nuclei exhibit an elongated shape, were also excluded. A metadata file was then created for each movie file containing the position as a fraction of the embryonic length (0 for anterior, 1 for posterior), the nuclear cycle (determined by counting the number of mitoses before the 14^th^ cycle). Visual examination of the dataset was used to determine if there was any motion of the nuclei during the acquisition period. Movies that contained any detectable motion were discarded or cropped to only include the time interval where there was no motion. A rectangular region of interest was then drawn around each nucleus which was then used to crop areas around individual nuclei. The boundary of each nucleus was then marked using a hand-drawn polygon. A masked movie was then created for each nucleus where regions outside the nucleus were set to 0 gray scale values so that all the analyses described below were only performed on molecules within nuclear regions.

### Single molecule localization and tracking using Dynamic Multiple-target tracing (MTT)

Localization and tracking of single molecules was conducting using a MATLAB implementation of the MTT algorithm (Serge, Bertaux, Rigneault, & Marguet, 2008). In brief, the algorithm first performs a bi-dimension Gaussian fitting to localize particles constrained by a log-likelihood ratio test subject to the localization error threshold. Deflation looping is performed to detect molecules that are partially overlapping. The parameters of the localization and tracking algorithms were empirically determined through iterative examination of the results. For all data sets the following settings were used: For localization, maximum number of deflation loops was set to 10, localization error to 10^−6^. For tracking the maximum expected diffusion coefficient was set to 5 μm^2^/s, maximum number of competitors was set to 1, and maximum off/blinking-frames was set to 1.

### Residence time analysis

The residence times were estimated from the 100 millisecond data using the results from the single particle tracking using the MTT algorithm. The data was pooled into bins corresponding to the position along the A-P axis of the embryo in 1/3 fractions of the embryonic length (EL, 0-1 anterior to posterior), with the following number of nuclei and single molecule trajectories per position bin:

Anterior: 34 nuclei, 17735, trajectories
Middle: 70 nuclei, 40092 trajectories
Posterior: 83 nuclei; 20823 trajectories

In the ZLD- embryos we measured the following number of nuclei and single molecule trajectories per position bin:

Anterior: 23 nuclei, 11415, trajectories
Middle: 31 nuclei, 7572 trajectories
Posterior: 31 nuclei; 3606 trajectories

The survival probability distribution was then calculated as 1- the cumulative distribution function of the trajectory lengths and was fit to both single and double exponential models(Mazza, Abernathy, Golob, Morisaki, & McNally, 2012). The double exponential model fit the data better in all cases (Figure 1-figure supplement. 3 and 4). The model used to fit the data and to calculate the time constants and fraction of the population is:

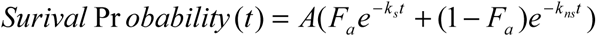

Where *k_s_* and *k_ns_* are the low (specific) and high (non-specific) off-rates respectively. The total pooled datasets of 78650 trajectories from the MTT results from 187 nuclei were also fit in the same manner.

To correct for photo-bleaching we used a *His2Av-eGFP* to estimate the bleaching constant (0.00426 s^-1^) and correct the off-rate as k_s,corrected_=k_s_-k_bleach_.(Table 1). We note that the bleaching correction has minimal effect on our estimated off-rates. The fact that we are not limited by bleaching due to the transient nature of BCD binding is further validated through even longer, 500 milliseconds exposure time measurements on 17 nuclei which provides an estimate for the specific and non-specific off-rates that do not differ significantly from those measured at 100 milliseconds (Figure 1-figure supplement 4 and Table 1). The results of the fits to all data are shown in Table 1.

### Fluorescence Recovery After Photobleaching (FRAP)

Experiments were performed on a Zeiss LSM 800 laser scanning confocal system (coupled to a Zeiss Axio Observer Z1) using a Plan-Achromat 63x/1.4 NA Oil immersion objective and GaAsP-PMT detector and a 488 nm laser. A circular bleach region with a diameter of 1.5 μm was used and the bleach location was selected manually in each nucleus at approximately the center and a total bleach time of 78.1 ms was used. Data was acquired for at least 1 second prior to bleaching and for at least 6 seconds post bleaching, with a time interval of 0.430 ms. Experiments were performed using live embryos from the same fly-line as for lattice light-sheet imaging and which were collected and prepared in the same manner as described above with the exception of the mounting procedure. For FRAP the embryos were mounted between a semipermeable membrane (Biofolie; In Vitro Systems & Services) and a coverslip and then embedded in Halocarbon 27 oil (Sigma). As in the case of the single molecule measurements, the embryos were staged using the HIS2-AV-MRFP channel and all data was acquired on embryos in nuclear cycle 13. The data was acquired on 21 nuclei all within the first 25% of the embryonic length from the anterior pole. FRAP experiments could not be performed at more anterior locations due to the low BCD concentrations and thus signal to noise ratio.

For analysis, the spatial and temporal location of the bleach region was retrieved from the meta-data recorded by the microscope control software and manually verified by inspecting the data. Due to the short duration of the movies and rapid recovery drift correction was not necessary. Each nucleus was manually segmented from the rest of the image by defining a polygon region of interest. A region of interest the same size as the bleach region was also marked in an area of each image outside of the nuclear region to be used to measure the background or dark intensity. The FRAP curve for each nucleus was then calculated as follows:

First, the mean intensity in the nuclear region, I_nuc_(t), bleach region, I_bleach_(t), and background region I_dark_(t) regions were calculated at each frame.

The background intensity was them subtracted from both the mean nuclear and bleach region intensities. To correct for photo-bleaching from imaging the bleach region intensity was then divided by the mean nuclear intensity at each time point. The resulting photo-bleaching corrected intensity was then normalized to the mean pre-bleach intensity, calculated as the average intensity in all frames prior to bleaching (I_pre-bleach_). The final FRAP curve for each nucleus was thus calculated as:

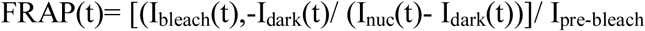

The calculated FRAP curves for each nucleus were then aligned to the bleach frame and averaged to generate the average FRAP curve shown in Fig. 1D. The averaged recovery data was then fit to both a single exponential (1-A*exp([-ka*t]) and double exponential (1-A*exp([-ka*t]- B*exp([-kb*t]) model (Figure 1-figure supplement 5 A,B), to measure the recovery time constants. There is no significant difference in the quality of fit between the two models. For comparison, the exact same experimental and analysis procedure was followed for His2AV-MRFP1 (Figure 1-figure supplement 5 C) in the same embryos with the exception of using a 561 nm bleach laser, 3 nuclei were measured with these settings. For the histone measurements the two-exponent fit was significantly better than the one-exponent as expected (Figure 1-figure supplement 5).

### Displacement Distribution Analysis

Single molecule trajectories were analyzed as described above. A total of 158 nuclei from 4 embryos were analyzed. The data from nuclei were binned according to their position along the A-P axis in 1/3 fractions of the embryonic length (EL, 0-1 anterior to posterior), with the following number of nuclei and single molecule displacements per bin: In the anterior most (0- 0.2) positions, tracking could not be performed reliably due to the high concentrations of Bicoid at those locations at these high frame rates and presumably a large mobile population.

Anterior: 30 nuclei, 12923, trajectories
Middle: 67 nuclei, 23640 trajectories
Posterior: 66 nuclei; 8600 trajectories

The fraction of the population that is bound vs. mobile was estimated using two approaches. First, a cumulative distribution function of the displacements was calculated for each EL bin (Figure 2-figure supplement 1A), displacements corresponding to distances less than a value of 225 nm were scored as part of the bound population. In the second approach the probability density functions of the displacements were fit to a two population model (Figure 2-figure supplement 1B).

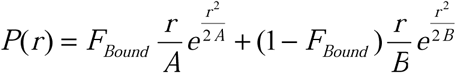

Where *F_Bound_* is the fraction of the population that is considered bound and *r* is the displacement distance. The two population model fits the displacement data with R^2^ values of 0.97, 0.98, and 0.96, for the anterior, middle and posterior distributions respectively and provides an estimate of the fraction present in the mobile and bound populations. The trend of an increase in the population bound estimate in the middle and posterior positions relative to the anterior is similar from both approaches. Although the bound population includes non-specific binding events as well, what we are interested in the relative change across the A-P axis.

We note that the diffusion coefficients for the bound and mobile populations can in principle be estimated from displacement distributions, however as explained by Mazza et al. (Mazza et al., 2012):, they cannot be estimated accurately from a fit to a displacement distribution at a single time step and are thus not reported. To accurately measure the diffusion coefficients more stable and photo switchable fluorophores are necessary to be able to track single molecules for more time points and at higher time resolutions.

### DBSCAN based analyses of clusters

The clustering of Bicoid binding events in the 100 millisecond WT embryos dataset is readily apparent in nuclei across the A-P axis in the projection of all localizations from the MTT algorithm (Figure. 2B). In order to automatically identify clusters and count the number of detections/ cluster vs. the whole nucleus (Figure. 2C), a MATLAB implementation (Yarpiz Team 2015) of the widely used Density-based spatial clustering of applications with noise (DBSCAN) was used with a minimum points setting of 10 points and an epsilon (maximum radius of neighborhood) setting of 0.8. These settings were empirically determined by iteratively changing parameters and examining the results. A balance had to be struck in an ability to accurately identify clusters in the high density situations in anterior nuclei and also low density situations in the posterior. Only datasets with timespans of at least 105 seconds were included. A total of 12, 49, and 48 nuclei and 436, 1168, and 367 clusters were analyzed in this manner for the Anterior, Middle, and Posterior positions respectively. Examples of clusters identified in nuclei at various positions are shown in Figure 3-figure supplement 2. For comparing distributions outliers were removed (< 5^th^ or > 95^th^ percentile) to disregard clusters that were significantly over or under-partitioned. In the case of the ZLD- embryos, due to the loss of apparent clustering, DBSCAN was not able to detect clusters so instead the spatial distribution of points was compared using pair-correlation analysis as described below to provide insight on the change beyond visual examination.

### Pair correlation analysis of clustering

The pair correlation function for the spatial distribution of particles computes the probability of finding a particle at the range of distances from another particles. In the case of complete spatial randomness which can be represented by a Poisson process the pair correlation function is equal to 1. The analysis is conducted on point lists generated from the MTT algorithm using the first spatial coordinate of each trajectory. A total of 22, 48, and 42 nuclei were used for the Anterior, Middle, and Posterior positions respectively for the WT embryos and a total of 23, 31, and 31 nuclei were used for the Anterior, Middle, and Posterior positions respectively for the *zld-* embryos. A MATLAB implementation (Ilya Valmianski 2014) was used to calculate the correlation function for the spatial distribution in each nucleus, the results were then averaged for each position (Anterior, Middle, Posterior) embryo for comparison (Figure 3-figure supplement 3).

### Chromatin Immunoprecipitation and Sequencing (ChIP-Seq)

Embryos were collected from a population cage for 90 minutes and then aged for 2 hours in order to enrich for embryos at developmental stage five. Embryos were then fixed with formaldehyde as previously described (Li et al., 2008) and sorted by morphology for those at early stage 5. The posterior thirds of embryos were sliced off by hand with a scalpel. A pool of the embryo segments from approximately 300 embryos was combined with 20 μg of whole *Drosophila pseudoobscura* embryos at stage 5 (to serve as carrier), and homogenized in homogenization buffer containing 15mM NaCl, 15mM TrisHCl pH 7.5, 60mM KCl, 1mM EDTA, and 0.1% Triton-X100, with 1mM DTT, 0.1 mM PMSF, and protease inhibit cocktail (Roche) added before use. After homogenization, 0.5% NP40 was added, and following a 5 minute incubation, samples were spun down at a low speed. Nuclei in the pellet were then washed once with the homogenization buffer containing 0.2M NaCl. The low speed centrifugation was repeated, and the recovered nuclei pellet was then re-suspended in nuclear lysis buffer (10 mM TrisCl, pH 7.9, 100 mM NaCl, 1 mM EDTA, 0.5 % Sarkosyl) +1%SDS, +1.5% sarkosyl. The chromatin was recovered by spinning the sample at full speed in a micro-centrifuge at 4^o^C for 1 hour and was re-suspended in a small volume of nuclear lysis buffer. Chromatin was sheared to an average size of 300bp using a Covaris sonicator (peak power= 140, duty factor = wo, cycle burst = 200, time = 2:20 minutes). Chromatin immunoprecipitation was performed using 72ng of chromatin and 1.5ug of an anti-Bicoid polyclonal antibody described previously (Li et al., 2008). The Bicoid antibody was coupled to Dynabead M-280 sheep anti-rabbit magnetic beads and the immunoprecipitation was conducted with the standard protocol from the manufacturer. DNA libraries for the chromatin immunoprecitation samples were prepared using the Rubicon genomics thru-plex DNA-seq kit using 16 PCR cycles and sequenced using Illumina High-seq with 2500 rapid run100bp single end reads. The sequencing reads were aligned to a combined *D. pseudoobscura* (Flybase Release 1.0) and *D. melanogaster* genome (Flybase Release 5) using Bowtie with options set as -3 70 -n 2 -m 1. The aligned reads were converted to wig files using custom scripts available upon request. Wig files were normalized to 10 million mapped reads.

### ChIP-Seq analysis

The posterior embryo ChIP-seq data was compared to published data on whole embryos from the same developmental stage (Bradley et al., 2010) downloaded from the NCBI GEO database with accession number GSM511083. To compare against Zelda binding, previously published data (Harrison et al., 2011) was downloaded from the NCBI GEO database with accession number GSM763061. Since the purpose of the analysis was to compare relative enrichment at genomic loci over each datasets respective background, the following normalization procedure was used: First, for each chromosome the average of the signal over the entire chromosome (a proxy for the background signal) was subtracted, negative values were then treated as below the background and discarded. The data was then normalized to the background subtracted average of each chromosome such that the normalized data now represents enrichment over background. For visualization data was smoothed using a Savitzky-Golay filter of order 3 over a 0.250 kbp sliding window. For analysis on CRMs the RedFly annotation database was used. For analysis centered on called peaks in either the whole embryo BCD data or the Zelda data .bed files containing peak locations were downloaded from the NCBI GEO database.

## ACKNOWLEDGMENTS

We thank Astou Tangara and the Betzig group at HHMI Janelia Research campus for help in constructing the Lattice Light Sheet Microscope. We thank Robert Tjian for extensive discussions and advice along the course of this work, and for his help in writing this manuscript. We thank Elizabeth Eck for her help in generating the zelda germ-line clones. We thank members of the Darzacq, Tjian, Garcia, and Eisen labs for suggestions and discussion. This work was supported by the California Institute of Regenerative Medicine (CIRM) LA1-08013 and the National Institutes of Health (NIH) UO1-EB021236 & U54-DK107980 to X.D., by the Burroughs Wellcome Fund Career Award at the Scientific Interface, the Sloan Research

Foundation, the Human Frontiers Science Program, the Searle Scholars Program, and the Shurl & Kay Curci Foundation to H.G., a Howard Hughes Medical Institute investigator award to M.E., J.H. and A.R are supported by NSF Graduate Research Fellowships.

## VIDEO CAPTIONS

**Video 1**-Movies corresponding to the still frames shown in Figure 1A.

**Video 2**- Representative data from a 90 second segment of a 100 millisecond exposure time movie acquired at an anterior position (EL (x/L) of 0.1). Top left shows the raw data and top right the corresponding surface plot representation. Bottom left shows a running max projection of the data and bottom right shows a surface plot representation of the same

**Video 3**-Representative data acquired at 10 millisecond exposure times for 4 nuclei.

**Figure 1-figure supplement 1.**
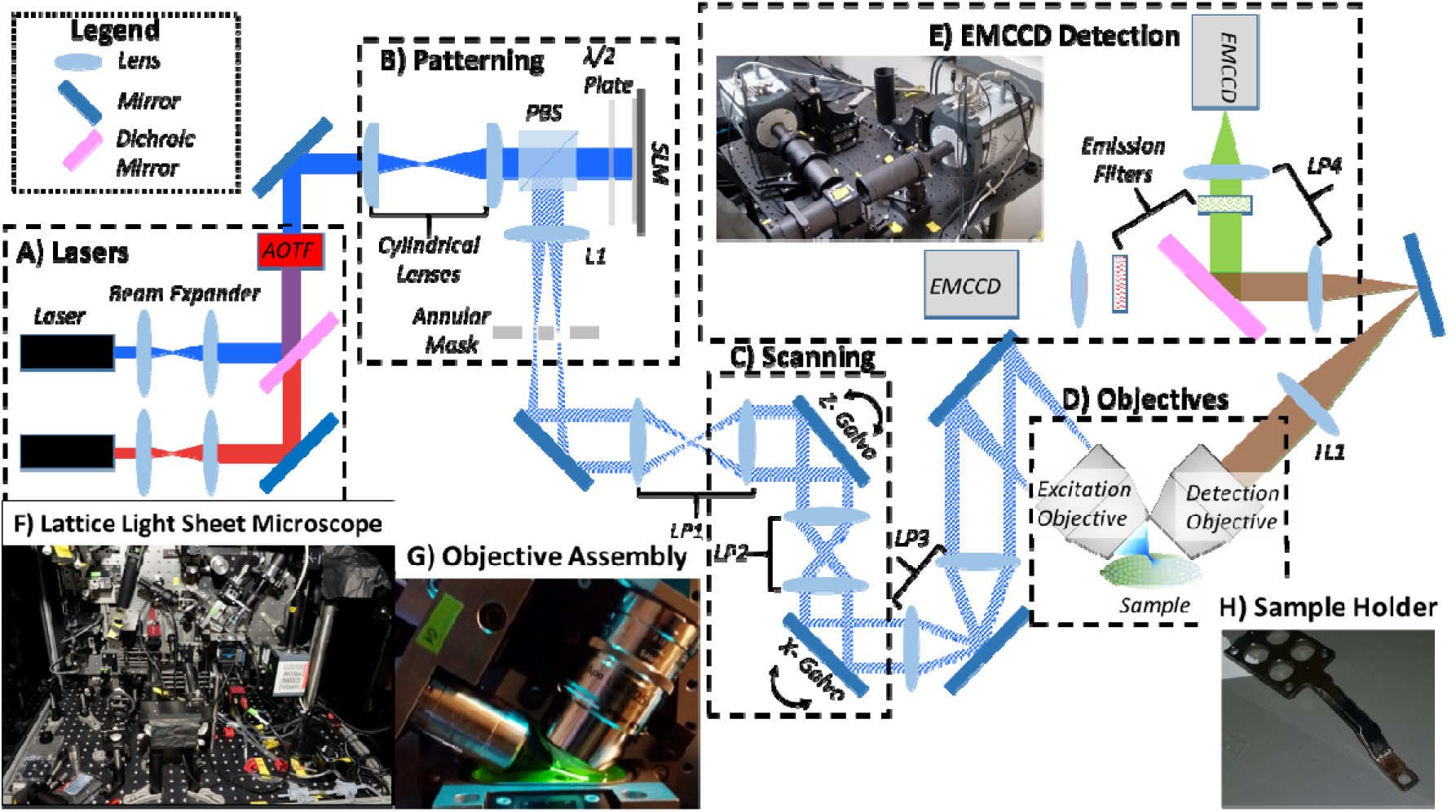
Lattice Light-Sheet Microscope Implementation. The Lattice light-sheet was built as described by Chen et al, Science, 2014. This simplified schematic shows the major modules of the microscope as follows: **(A)** Laser Module, contains 6 lasers ranging from 405 nm to 639 nm, which are independently expanded, collimated, and input into an Acousto-Optic Tunable Filter (AOTF). **(B)** The patterning module contains a pair of cylindrical lens to expand the input Gaussian beam to a stripe, and a half-wave (λ/2) plate, a polarizing beam splitter (PBS), and a Spatial Light Modulator (SLM) to perform the patterning.Lens L1 projects the Fourier Transform of the SLM pattern onto an annular mask for spatial filtering (**C**) Lens pair 1 (LP1) is then used to de-magnify the annular mask plane and project it onto the z-scan galvo and LP2 projects the z-galvo plane onto the x-scanning galvo. (**D**) LP3 magnifies the x-galvo plane and projects it onto the back pupil plane of the excitation objective, where it is focused (Fourier transform) to project the final light sheet pattern onto the sample. Emitted fluorescence is collected by the detection objective. (**E**) The tube lens (TL1) focuses an intermediate image plane which is then magnified using LP4 and projected onto an EMCCD. **A** dichroic mirror is placed between the first and second lenses of LP4 to split the signal into high and low wavelengths. An emission filter is placed after the dichroic on both sides to select the chromatic bandwidth of the signal reaching each EMMCD independently. Inset shows an image of the EMCCD detection module (**F**) Image of our lattice light sheet microscope. (**G**) Image of the Objective assembly with a dye solution in the sample chamber to visualize the excitation light. (**H**) Image of the sample holder on which the 5 mm coverslip is mounted.

**Figure 1-figure supplement 2.**
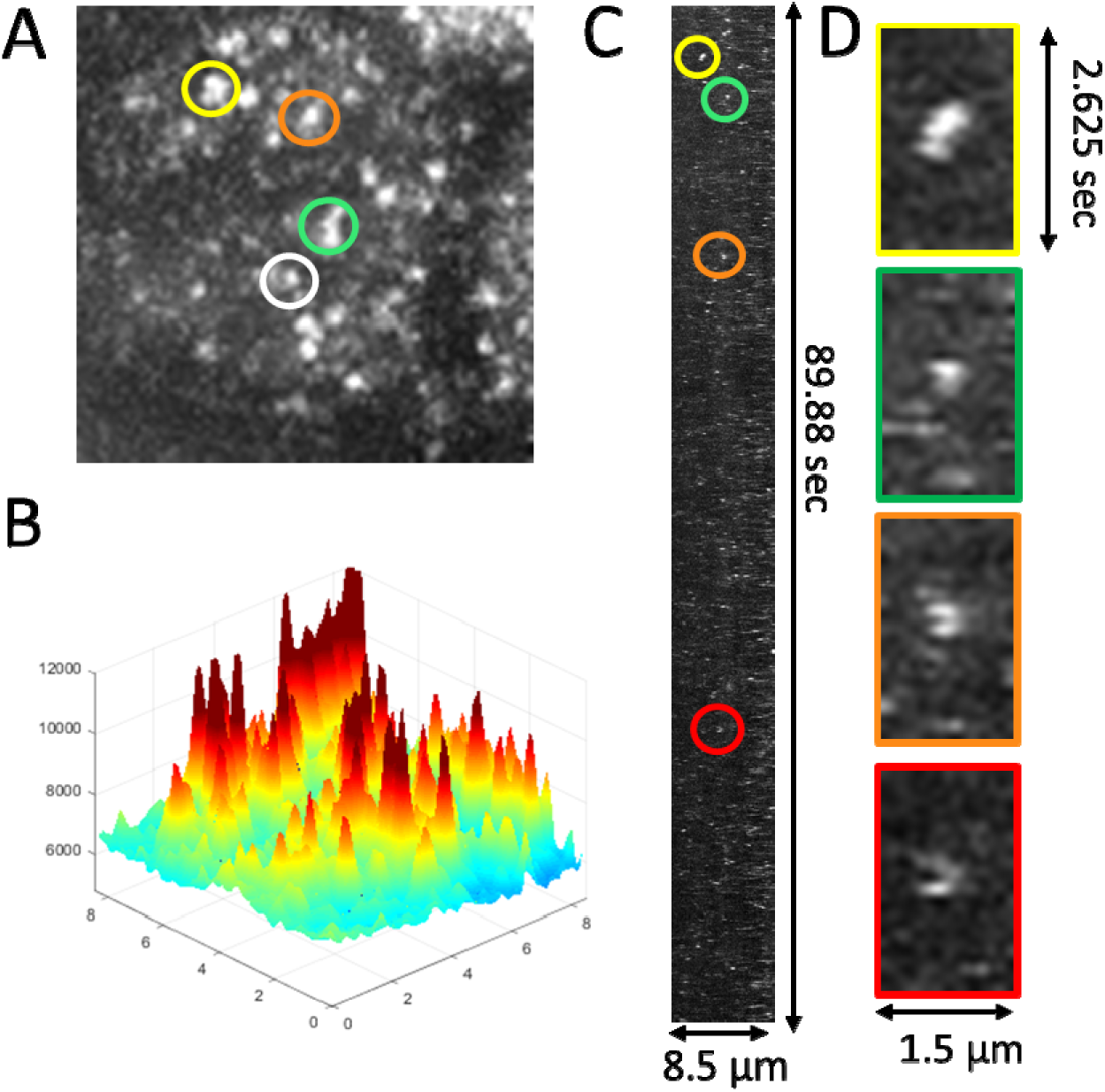
Single Molecule Imaging of BCD-eGFP at 100 milliseconds to estimate residence times. **(A)** The max projection in time of a 90 second segment of a representative 100 millisecond dataset acquired at an anterior nucleus (EL (x/L) of 0.1), corresponding to the last frame of Video 2. **(B)** Surface plot representation of (A) to illustrate the signal-to-background ratio of single molecule binding events. The smaller peaks likely correspond to slowly diffusing molecules that are not in the imaging volume for the entire exposure time. **(C)** Maximum projection through x-t (kymograph representation) of the data shown in Video 2 from which (A-B) were calculated. Colored circles correspond to those in (A). **(D)** Zoomed in x-t view of the circled regions in (A) and (C), illustrating the transient nature of BCD binding.

**Figure 1-figure supplement 3.**
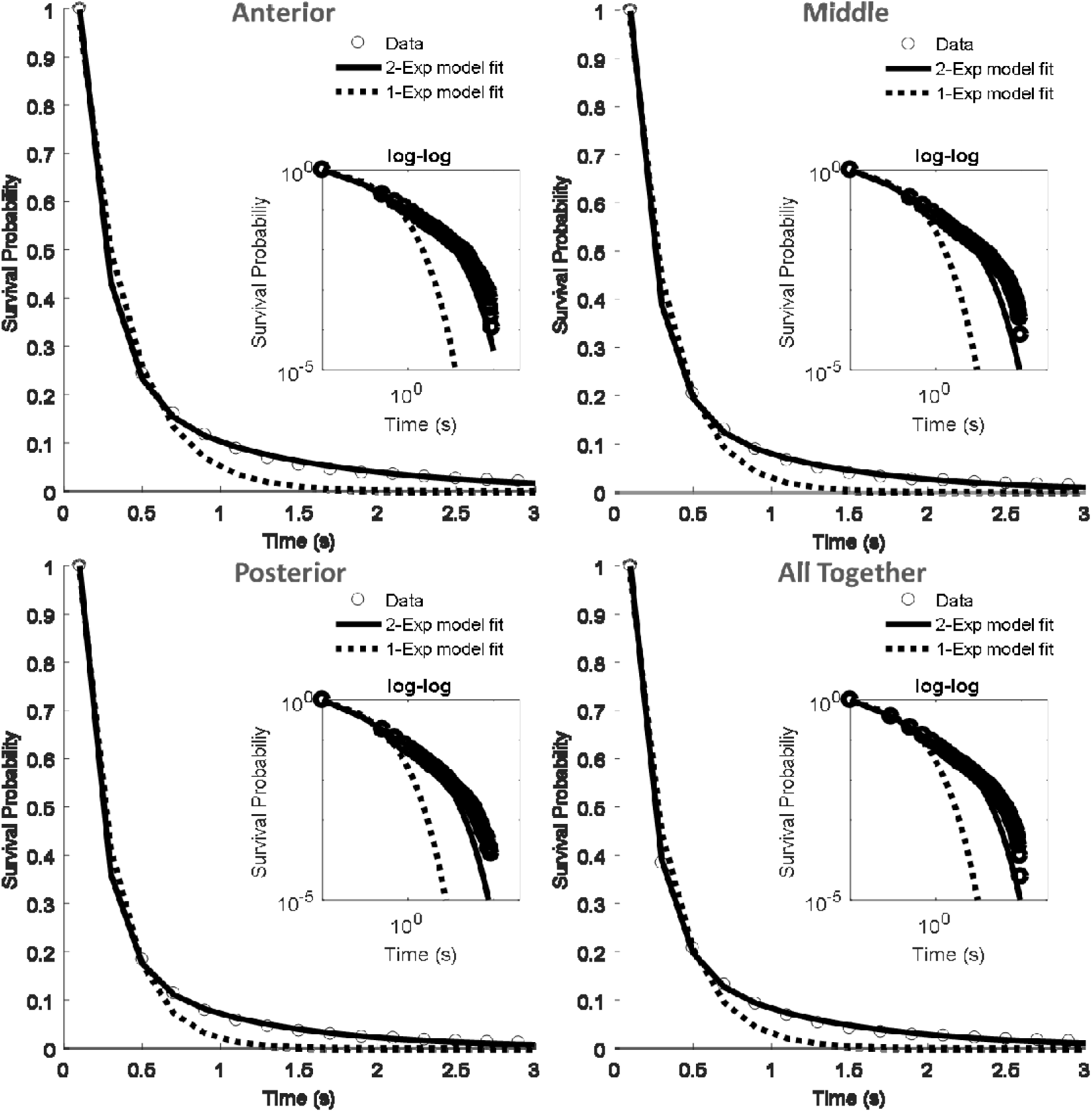
Fits to the survival probability distributions of the 100 millisecond datasets. The survival probability distributions calculated from the trajectories output by the MTT analysis on the 100 millisecond data set for the Anterior (34 nuclei, 17735,trajectories), Middle (70 nuclei, 40092 trajectories), Posterior (83 nuclei; 20823 trajectories) segments and all the data pooled together (187 nuclei, 78650 trajectories). The solid lines show the fit to a 2 exponent-model and the dashed lines to a 1-exponent model. Insets show the same on a log-log scale to visualize the difference between 1 and 2 exponent fits at longer time scales. All two-exponent fits have an R^2^ value>0.99. See Table 1 for a summary of the fit parameters.

**Figure 1-figure supplement 4.**
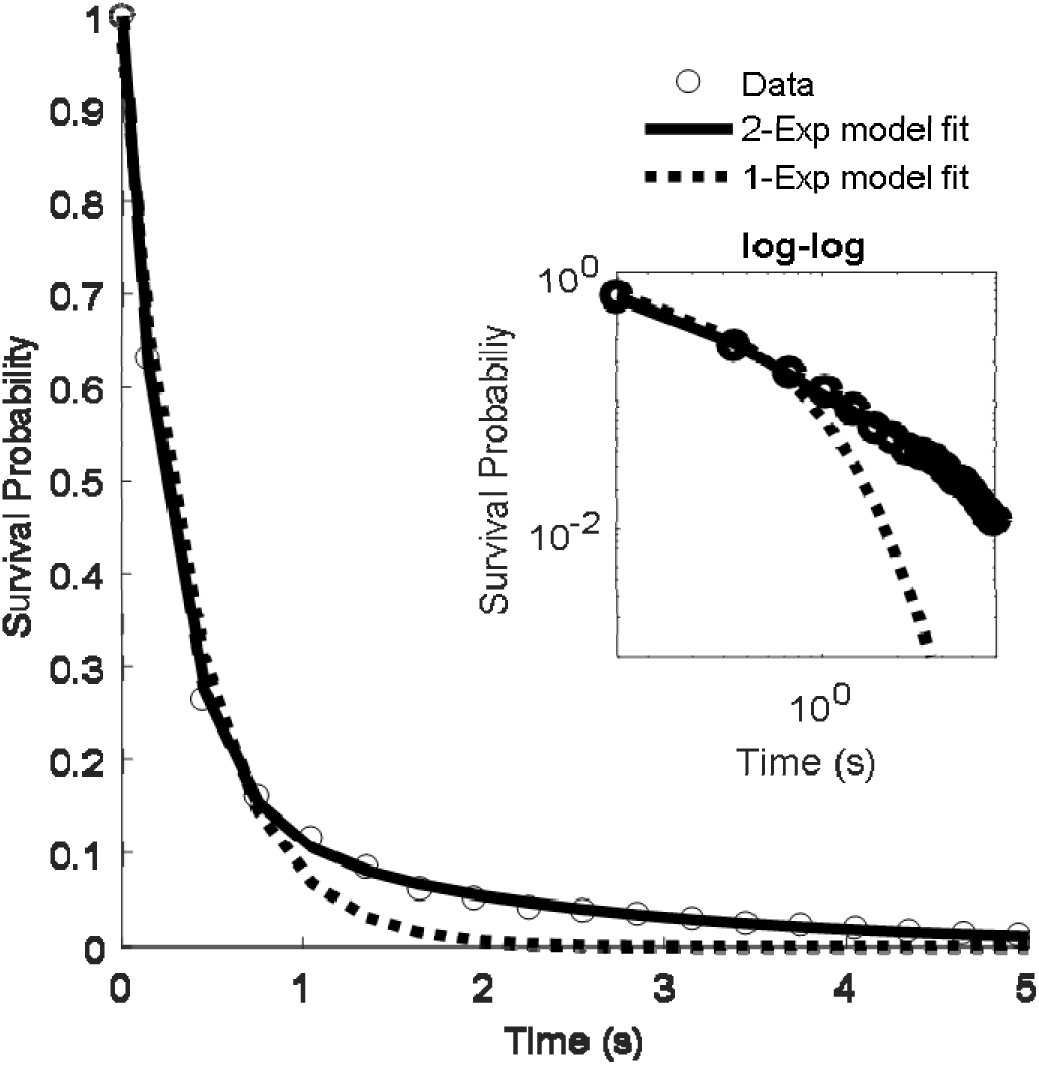
Fits to the survival probability distribution of the 500 millisecond dataset. The survival probability distribution of residence times calculated from the MTT algorithm on the 500 ms data set of 17 nuclei and 1211 trajectories. The solid lines show the fit to a 2-exponent model and the dotted lines to a 1-exponent model. The insets shows the same on a log-log scale to visualize the difference between 1 and 2 exponent fits. R^2^ value for the 2-exponent fit is 0.99

**Figure 1-figure supplement 5.**
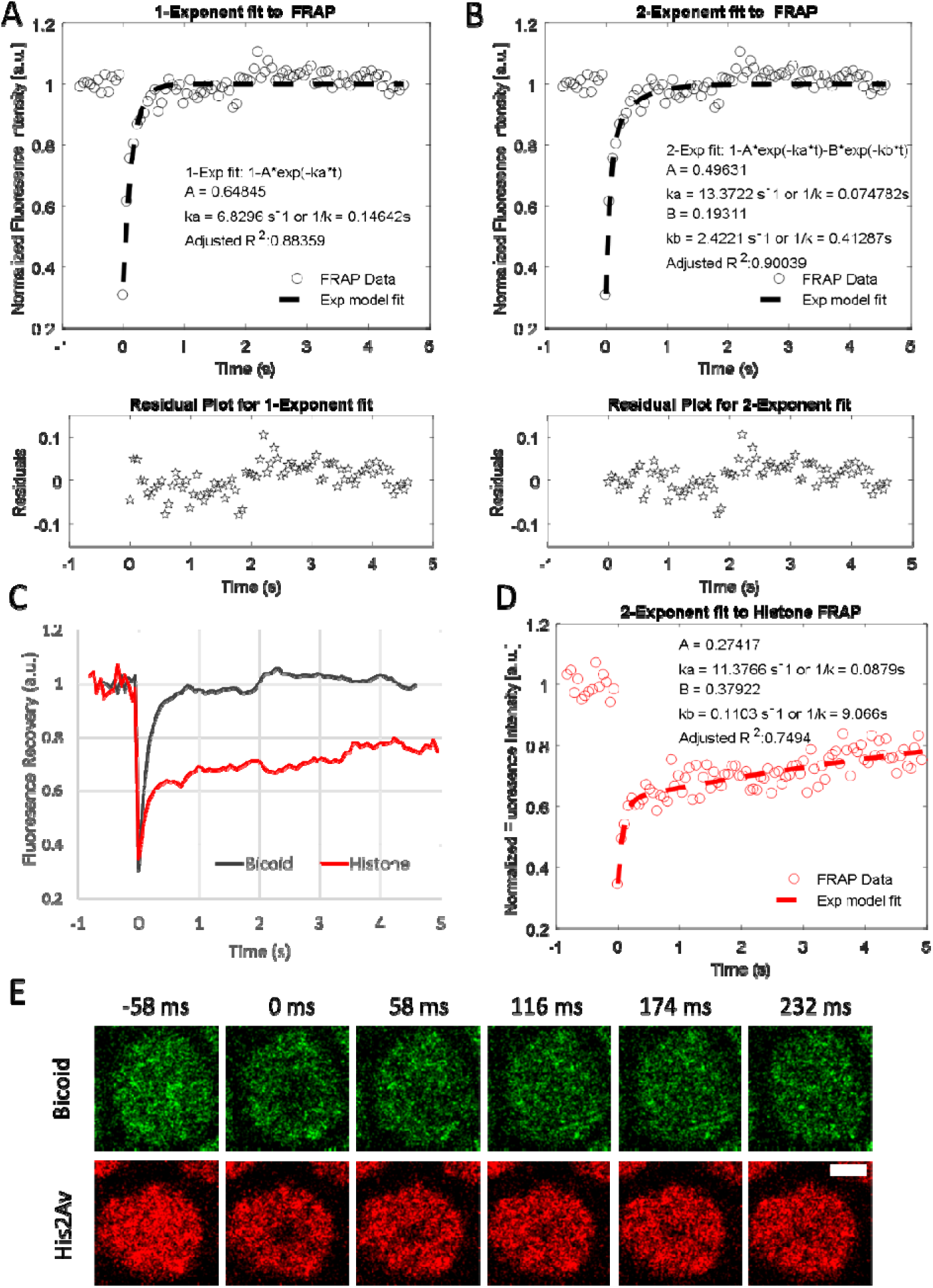
Analysis of FRAP data. **(A)** Averaged Bcd-eGFP FRAP data and single exponential fit results. **(B)** Double exponential fit results to average Bcd-eGFP FRAP. **(C)** Comparison of averaged BCD-eGFP (21 nuclei) and His2AV FRAP data (3 nuclei). **(D)** Double exponential fit to Histone (His2AV) FRAP. **(E)** Representative images from BCD-eGFP and His2AV FRAP experiments, white scale bar is 2μm.

**Figure 2-figure supplement 1.**
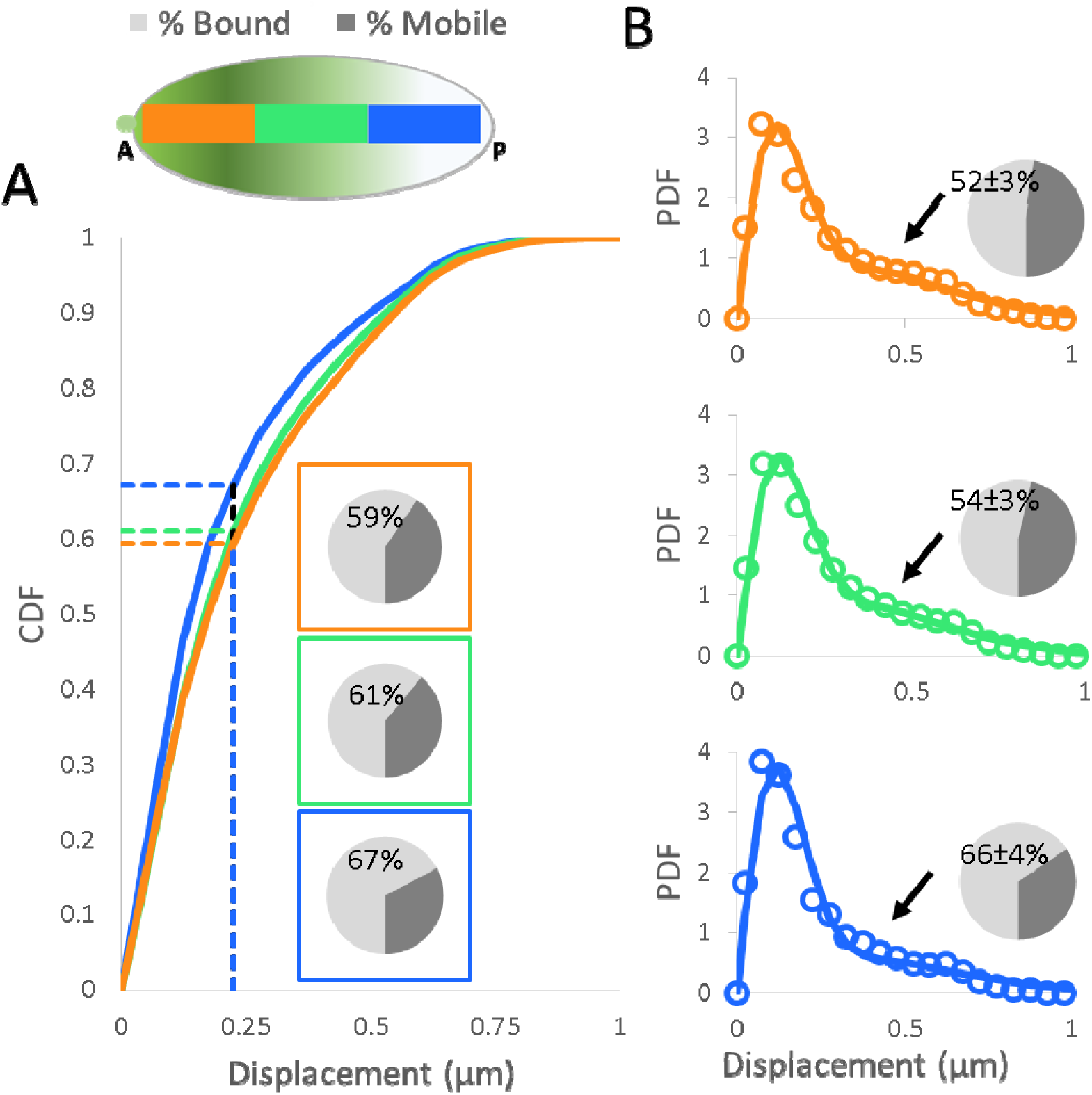
Analysis of Displacement Distributions. (A) Cumulative distribution functions of the displacement data, pie charts indicate the Bound and Mobile fraction as measured by the CDF value at 0.225 μm (dashed lines). (B) Probability Density Function of the displacement data (markers) and fits to the two population model (solid lines), pie charts show the mobile and bound fractions from the results of the fit, labels indicate the bound fraction and the error is the standard error associated with the fit parameter. Arrows point to the mobile population in the distributions that is decreasing from anterior to posterior positions. (A-B) For each position the following number of nuclei and trajectories were included in the analysis Anterior: 30 nuclei, 12923, trajectories, Middle: 67 nuclei, 23640 trajectories, Posterior: 66 nuclei; 8600 trajectories.

**Figure 2-figure supplement 2.**
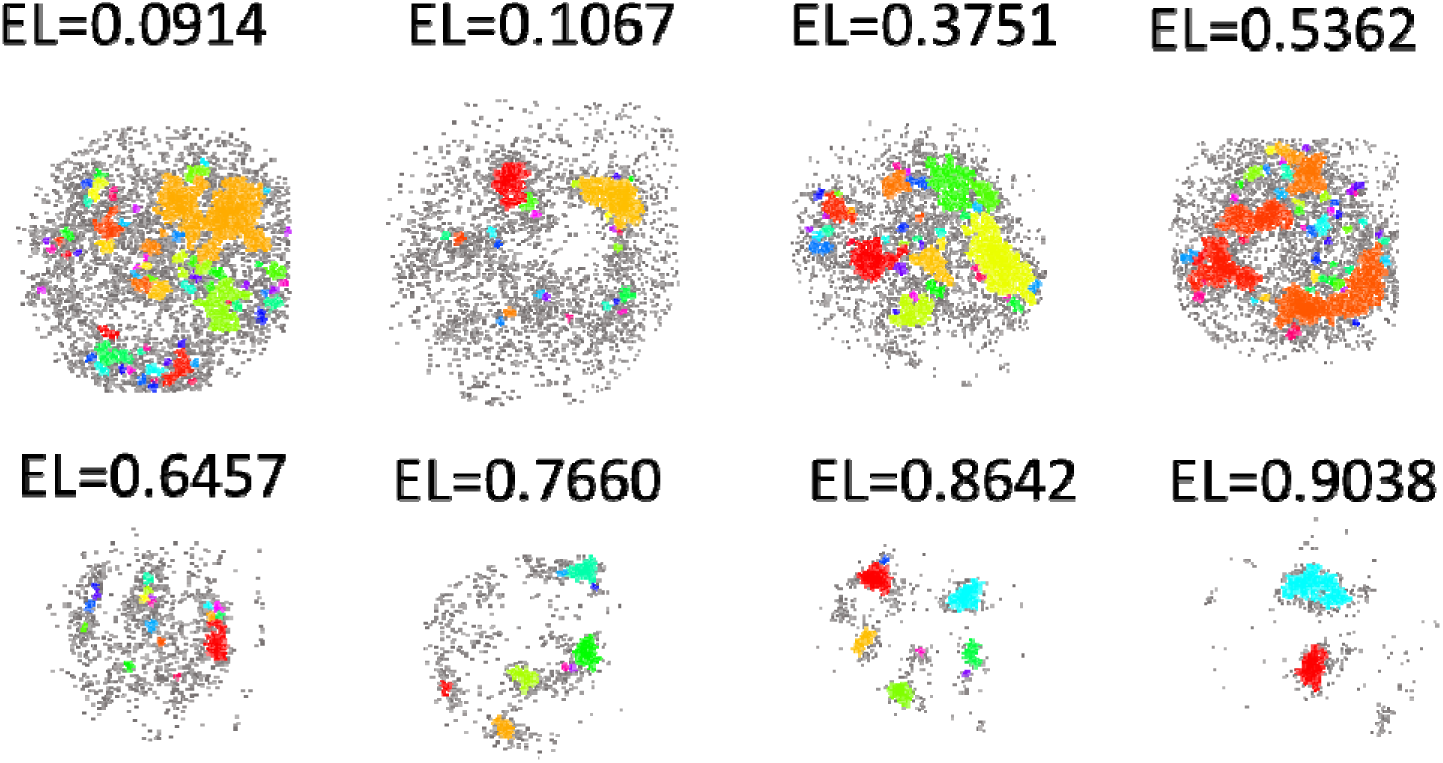
Cluster identification results from DBSCAN across the A-P axis. Examples of clusters identified along the A-P axis using DBSCAN with the position shown as fraction of embryonic length (EL). Particles included in the same clusters are represented with the same color.

**Figure 3-figure supplement 1.**
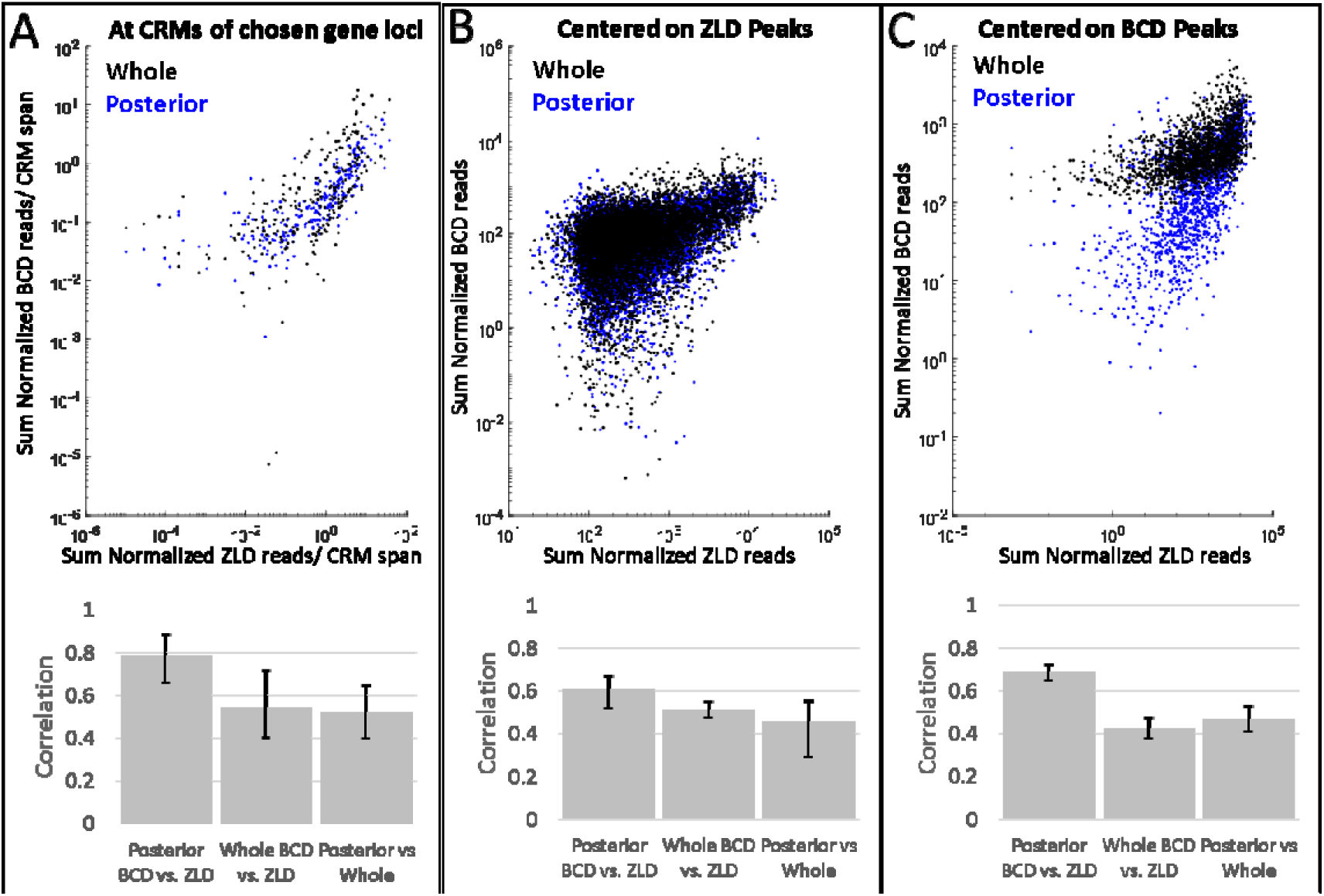
BCD binding in whole and posterior thirds embryos compared to ZLD binding. **(A)** Sum of Normalized BCD ChIP-seq reads divided by the length of the respective CRMs for posterior thirds (blue) and whole embryo (black) vs. Sum of Normalized ZLD ChIP-seq reads divided by the length of the respective CRM at annotated cis-regulatory modules (CRMs) of *eve*, *giant*, *hunchback*, *knirps*, *hairy*, *kruppel*, *caudal*, *fushitarazu*, *engrailed*, *wingless*, *runt*, and *gooseberry* loci. A total of 293 CRMs from the RedFly database were analyzed. **(A-B)** Sum of Normalized BCD ChIP-seq reads for posterior thirds (blue) and whole embryo (black) vs. Sum of Normalized ZLD ChIP-seq reads over a 500 bp region centered on **(B)** 8331 ZLD peaks and **(C)** 2145 BCD peaks. **(A-C)** Corresponding bar plots show Pearson correlation coefficients and error bars show 95% confidence intervals as determined by bootstrapping.

**Figure 3-figure supplement 2.**
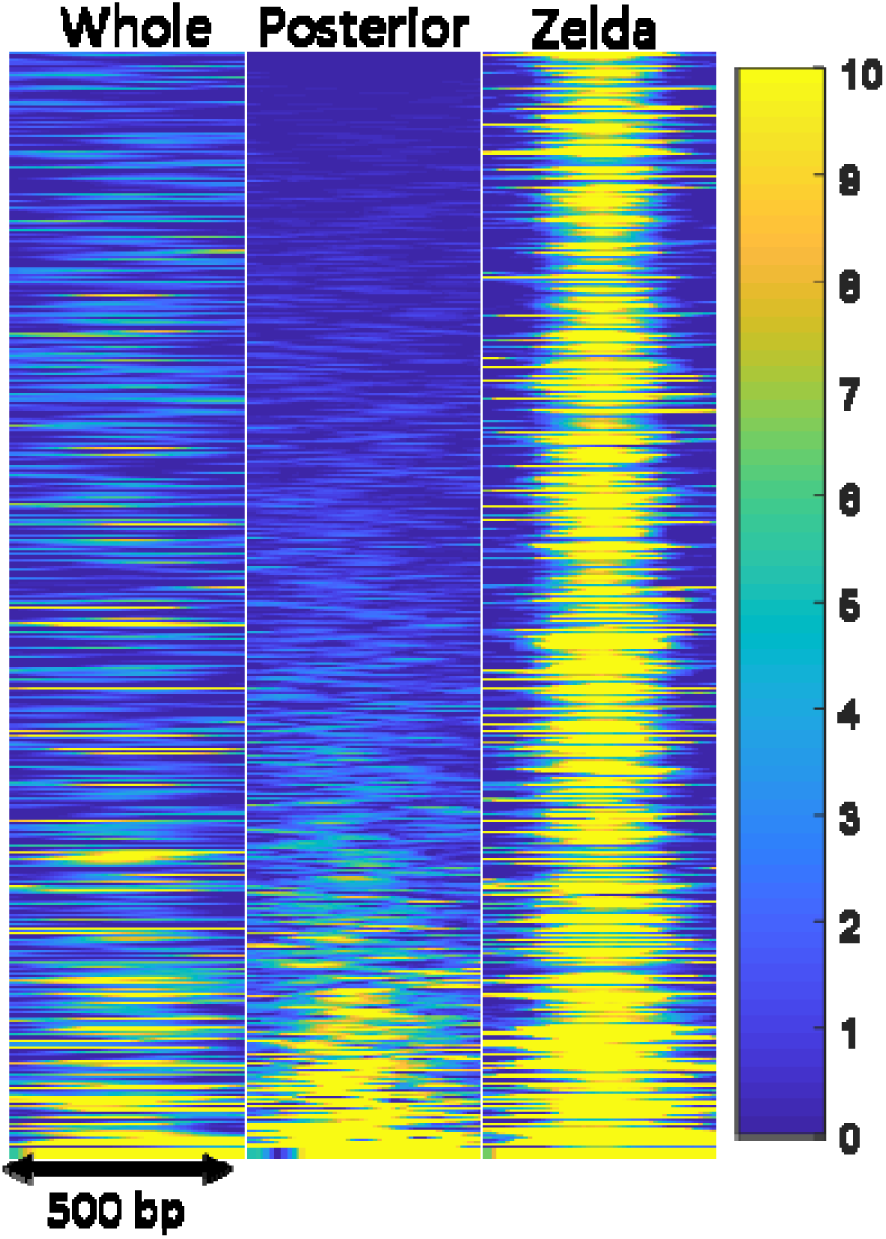
BCD and ZLD binding at Zelda peaks. Heat-map representation of normalized ChIP-seq reads for BCD (1^st^ two panels) and ZLD (3^rd^ panel) in a 500 bp window centered on ZLD peaks sorted according to increasing BCD signal in the posterior embryo data. A total of 8331 peaks are shown.

**Figure 3-figure supplement 3.**
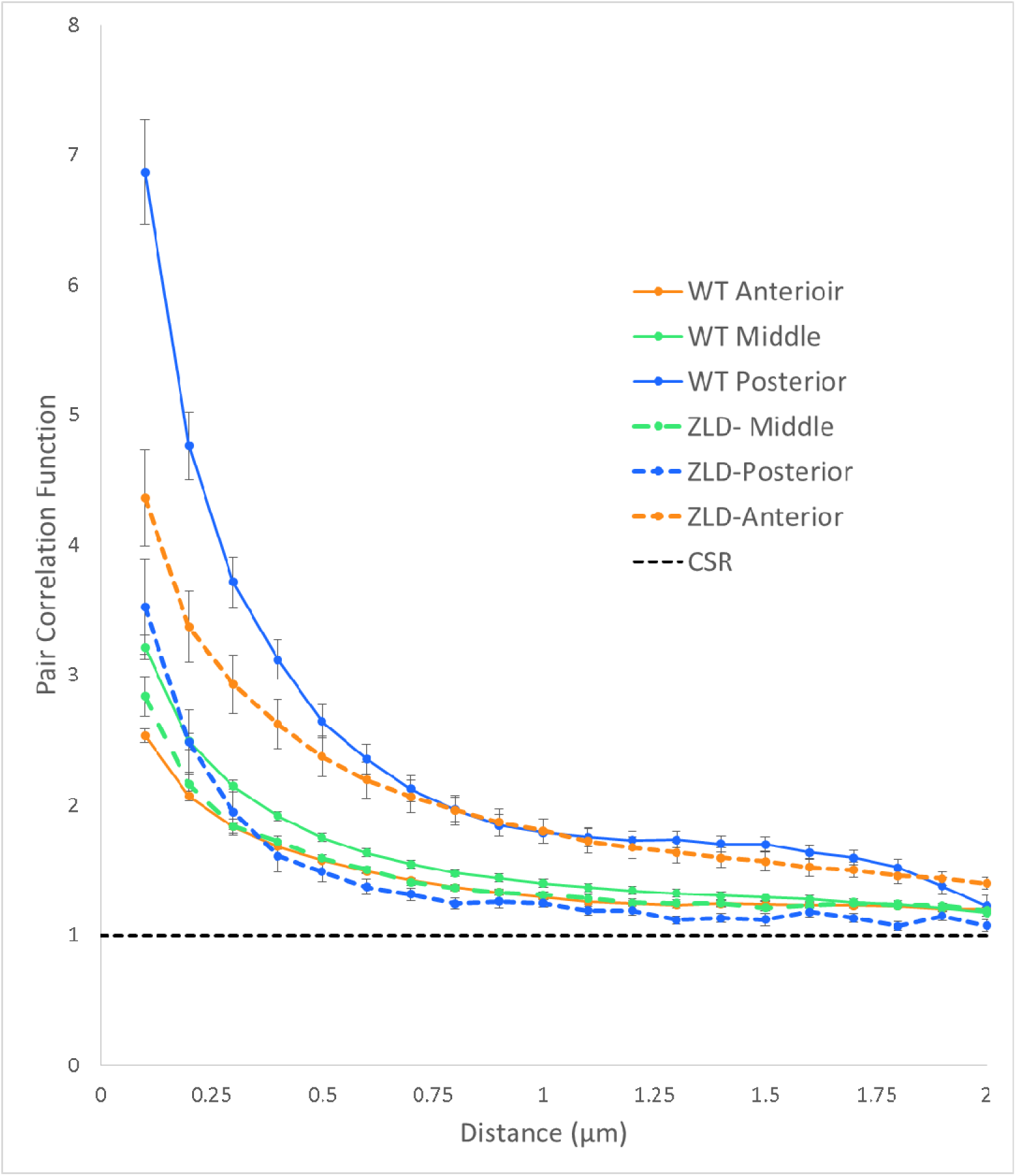
Averaged pair-correlation (radial distribution) functions for nuclei in the WT and ZLD- cases. Black dashed line indicates complete spatial randomness (CSR). Error bars indicate the standard error.

## REFERENCES

Bradley, R. K., Li, X. Y., Trapnell, C., Davidson, S., Pachter, L., Chu, H. C., … Eisen M. B. (2010). Binding Site Turnover Produces Pervasive Quantitative Changes in Transcription Factor Binding between Closely Related Drosophila Species. Plos Biology, 8(3). doi:ARTN e100034310.1371/journal.pbio.1000343

Burz, D. S., Rivera-Pomar, R., Jackle, H., & Hanes, S. D. (1998). Cooperative DNA-binding by Bicoid provides a mechanism for threshold-dependent gene activation in the Drosophila embryo. EMBO J, 17(20), 5998-6009. doi: 10.1093/emboj/17.20.5998

Chen, B. C., Legant, W. R., Wang, K., Shao, L., Milkie, D. E., Davidson, M. W., … Betzig, E. (2014). Lattice light-sheet microscopy: Imaging molecules to embryos at high spatiotemporal resolution. Science, 346(6208), 439-439. doi: DOI 10.1126/science. 1257998

Chen, H., Xu, Z., Mei, C., Yu, D., & Small, S. (2012). A system of repressor gradients spatially organizes the boundaries of Bicoid-dependent target genes. Cell, 149(3), 618-629. doi: 10.1016/j.cell.2012.03.018

Chen, J. J., Zhang, Z. J., Li, L., Chen, B. C., Revyakin, A., Hajj, B., …Liu, Z. (2014). Single-Molecule Dynamics of Enhanceosome Assembly in Embryonic Stem Cells. Cell, 156(6), 1274-1285. doi: DOI 10.1016/j.cell.2014.01.062

Cisse, II, Izeddin, I., Causse, S. Z., Boudarene, L., Senecal, A., Muresan, L., … Darzacq, X. (2013). Real-time dynamics of RNA polymerase II clustering in live human cells. Science, 341(6146), 664-667. doi: 10. 1 126/science. 1239053

Combs, P., & Eisen, M. (2017). Genome-wide measurement of spatial expression in patterning mutants of Drosophila melanogaster [version 1; referees: 2 approved] (Vol. 6).

Combs, P. A., & Eisen, M. B. (2013). Sequencing mRNA from Cryo-Sliced Drosophila Embryos to Determine Genome-Wide Spatial Patterns of Gene Expression. PLoS One, 8(8). doi: ARTN e71820 10.1371/journal.pone.0071820

Crocker, J., Noon, E. P., & Stern, D. L. (2016). The Soft Touch: Low-Affinity Transcription Factor Binding Sites in Development and Evolution. Curr Top Dev Biol, 117, 455-469. doi:10.1016/bs.ctdb.2015.11.018

Crocker, J., Tsai, A., Muthusamy, A. K., Lavis, L. D., Singer, R. H., & Stern, D. L. (2017). Nuclear Microenvironments Modulate Transcription From Low-Affinity Enhancers. bioRxiv. doi: 10.1101/128280

Driever, W., & Nusslein-Volhard, C. (1988a). The bicoid protein determines position in the Drosophila embryo in a concentration-dependent manner. Cell, 54(1), 95-104.

Driever, W., & Nusslein-Volhard, C. (1988b). A gradient of bicoid protein in Drosophila embryos. Cell, 54(1), 83-93.

Driever, W., & Nussleinvolhard, C. (1989). The Bicoid Protein Is a Positive Regulator of Hunchback Transcription in the Early Drosophila Embryo. Nature, 337(6203), 138-143. doi: Doi 10.1038/337138a0

Driever, W., Siegel, V., & Nusslein-Volhard, C. (1990). Autonomous determination of anterior structures in the early Drosophila embryo by the bicoid morphogen. Development, 109(4), 811-820.

Ephrussi, A., & St Johnston, D. (2004). Seeing is believing: the bicoid morphogen gradient matures. Cell, 116(2), 143-152.

Foo, S. M., Sun, Y., Lim, B., Ziukaite, R., O’Brien, K., Nien, C. Y., … Rushlow C. A. (2014). Zelda potentiates morphogen activity by increasing chromatin accessibility. Current Biology, 24(12), 1341-1346. doi:10.1016/j.cub.2014.04.032

Gregor, T., Wieschaus, E. F., McGregor, A. P., Bialek, W., & Tank, D. W. (2007). Stability and nuclear dynamics of the bicoid morphogen gradient. Cell, 130(1), 141-152. doi:10.1016/j.cell.2007.05.026

Hamm, D. C., Bondra, E. R., & Harrison, M. M. (2015). Transcriptional activation is a conserved feature of the early embryonic factor Zelda that requires a cluster of four zinc fingers for DNA binding and a low-complexity activation domain. Journal of Biological Chemistry, 290(6), 3508-3518. doi:10.1074/jbc.M114.602292

Hannon, C. E., Blythe, S. A., & Wieschaus, E. F. (2017). Concentration Dependent Chromatin States Induced by the Bicoid Morphogen Gradient. bioRxiv. doi: 10.1101/133348

Hansen, A. S., Pustova, I., Cattoglio, C., Tjian, R., & Darzacq, X. (2017). CTCF and cohesin regulate chromatin loop stability with distinct dynamics. Elife, 6, e25776. doi: 10.7554/eLife.25776

Harrison, M. M., Li, X. Y., Kaplan, T., Botchan, M. R., & Eisen, M. B. (2011). Zelda binding in the early Drosophila melanogaster embryo marks regions subsequently activated at the maternal-to-zygotic transition. PLoS Genet, 7(10), e1002266. doi: 10.1371/journal.pgen.1002266

La Rosee, A., Hader, T., Taubert, H., Rivera-Pomar, R., & Jackle, H. (1997). Mechanism and Bicoid-dependent control of hairy stripe 7 expression in the posterior region of the Drosophila embryo. EMBO J, 16(14), 4403-4411. doi: 10.1093/emboj/16.14.4403

Lebrecht, D., Foehr, M., Smith, E., Lopes, F. J. P., Vanario-Alonso, C. E., Reinitz, J., … Hanes S. D. (2005). Bicoid cooperative DNA binding is critical for embryonic patterning in Drosophila. Proceedings of the National Academy of Sciences of the United States of America, 102(37), 13176-13181. doi:10.1073/pnas.0506462102

Li, X. Y., Harrison, M. M., Villata, J. E., Kaplan, T., & Eisen, M. B. (2014). Establishment of regions of genomic activity during the Drosophila maternal to zygotic transition. Elife, 3. doi: ARTN e03737 10.7554/eLife.03737

Li, X. Y., MacArthur, S., Bourgon, R., Nix, D., Pollard, D. A., Iyer, V. N., … Biggin M. D. (2008). Transcription factors bind thousands of active and inactive regions in the Drosophila blastoderm. Plos Biology, 6(2), 365-388. doi:ARTNe2710.1371/journal.pbio.0060027

Liang, H. L., Nien, C. Y., Liu, H. Y., Metzstein, M. M., Kirov, N., & Rushlow, C. (2008). The zinc-finger protein Zelda is a key activator of the early zygotic genome in Drosophila. Nature, 456(7220), 400-U467. doi: 10.1038/nature07388

Liu, Z., Lavis, L. D., & Betzig, E. (2015). Imaging Live-Cell Dynamics and Structure at the Single-Molecule Level. Mol Cell, 58(4), 644-659. doi: 10.1016/j.molcel.2015.02.033

Liu, Z., Legant, W. R., Chen, B. C., Li, L., Grimm, J. B., Lavis, L. D., … Tjian R. (2014). 3D imaging of Sox2 enhancer clusters in embryonic stem cells. Elife, 3. doi: ARTN e04236 DOI 10.7554/eLife.04236.001

Lucchetta, E. M., Lee, J. H., Fu, L. A., Patel, N. H., & Ismagilov, R. F. (2005). Dynamics of Drosophila embryonic patterning network perturbed in space and time using microfluidics. Nature, 434(7037), 1134-1138. doi: 10.1038/nature03509

Mazza, D., Abernathy, A., Golob, N., Morisaki, T., & McNally, J. G. (2012). A benchmark for chromatin binding measurements in live cells. Nucleic Acids Res, 40(15), e119. doi:10.1093/nar/gks701

Morrison, A. H., Scheeler, M., Dubuis, J., & Gregor, T. (2012). Quantifying the Bicoid morphogen gradient in living fly embryos. Cold Spring Harb Protoc, 2012(4), 398-406. doi: 10.1101/pdb.top068536

Normanno, D., Boudarene, L., Dugast-Darzacq, C., Chen, J., Richter, C., Proux, F., … Dahan M. (2015). Probing the target search of DNA-binding proteins in mammalian cells using TetR as model searcher. Nat Commun, 6, 7357. doi: 10.1038/ncomms8357

Ochoa-Espinosa, A., Yu, D. Y., Tsirigos, A., Struffi, P., & Small, S. (2009). Anterior-posterior positional information in the absence of a strong Bicoid gradient. Proceedings of the National Academy of Sciences of the United States of America, 106(10), 3823-3828. doi:10.1073/pnas.0807878105

Ochoa-Espinosa, A., Yucel, G., Kaplan, L., Pare, A., Pura, N., Oberstein, A., … Small S. (2005). The role of binding site cluster strength in Bicoid-dependent patterning in Drosophila. Proceedings of the National Academy of Sciences of the United States of America, 102(14), 4960-4965. doi: 10. 1073/pnas.0500373102

Perry, M. W., Bothma, J. P., Luu, R. D., & Levine, M. (2012). Precision of hunchback expression in the Drosophila embryo. Current Biology, 22(23), 2247-2252. doi: 10.1016/j.cub.2012.09.051

Porcher, A., Abu-Arish, A., Huart, S., Roelens, B., Fradin, C., & Dostatni, N. (2010). The time to measure positional information: maternal Hunchback is required for the synchrony of the Bicoid transcriptional response at the onset of zygotic transcription. Development, 137(16), 2795-2804. doi:10.1242/dev.051300

Rivera-Pomar, R., Lu, X., Perrimon, N., Taubert, H., & Jackle, H. (1995). Activation of posterior gap gene expression in the Drosophila blastoderm. Nature, 376(6537), 253-256. doi: 10.1038/376253a0

Schulz, K. N., Bondra, E. R., Moshe, A., Villalta, J. E., Lieb, J. D., Kaplan, T., … Harrison M. M. (2015). Zelda is differentially required for chromatin accessibility, transcription factor binding, and gene expression in the early Drosophila embryo. Genome Research, 25(11), 1715-1726. doi: 10.1101/gr.192682.115

Serge, A., Bertaux, N., Rigneault, H., & Marguet, D. (2008). Dynamic multiple-target tracing to probe spatiotemporal cartography of cell membranes. Nat Methods, 5(8), 687-694. doi:10.1038/nmeth.1233

Small, S., Blair, A., & Levine, M. (1996). Regulation of two pair-rule stripes by a single enhancer in the Drosophila embryo. Developmental Biology, 175(2), 314-324. doi: DOI 10.1006/dbio.1996.0117

Struhl, G., Struhl, K., & Macdonald, P. M. (1989). The Gradient Morphogen Bicoid Is a Concentration-Dependent Transcriptional Activator. Cell, 57(7), 1259-1273. doi: Doi 10.1016/0092-8674(89)90062-7

Sun, Y. J., Nien, C. Y., Chen, K., Liu, H. Y., Johnston, J., Zeitlinger, J., & Rushlow, C. (2015). Zelda overcomes the high intrinsic nucleosome barrier at enhancers during Drosophila zygotic genome activation. Genome Research, 25(11), 1703-1714. doi: 10.1101/gr.192542.115

Tokunaga, M., Imamoto, N., & Sakata-Sogawa, K. (2008). Highly inclined thin illumination enables clear single-molecule imaging in cells. Nature Methods, 5(2), 159-161. doi: 10.1038/Nmeth.1171

Turing, A. M. (1952). The Chemical Basis of Morphogenesis. Philosophical Transactions of the Royal Society of London Series B-Biological Sciences, 237(641), 37-72. doi:DOI 10.1098/rstb.1952.0012

Wolpert, L. (1969). Positional Information and Pattern of Cellular Differentiation. Biophysical Journal, 9, A8-&.

Xu, H., Sepulveda, L. A., Figard, L., Sokac, A. M., & Golding, I. (2015). Combining protein and mRNA quantification to decipher transcriptional regulation. Nat Methods, 12(8), 739-742. doi: 10.1038/nmeth.3446

Xu, Z., Chen, H., Ling, J., Yu, D., Struffi, P., & Small, S. (2014). Impacts of the ubiquitous factor Zelda on Bicoid-dependent DNA binding and transcription in Drosophila. Genes Dev, 28(6), 608-621. doi:10.1101 /gad.234534.113

